# Machine learning-aided multidimensional phenotyping of Parkinson’s disease patient stem cell-derived midbrain dopaminergic neurons

**DOI:** 10.1101/2022.03.01.482490

**Authors:** Aurore Vuidel, Loïc Cousin, Beatrice Weykopf, Simone Haupt, Zahra Hanifehlou, Nicolas Wiest-Daesslé, Michaela Segschneider, Michael Peitz, Arnaud Ogier, Laurent Brino, Oliver Brüstle, Peter Sommer, Johannes H. Wilbertz

**Author notes:** Co-first authors (Aurore Vuidel, Loïc Cousin).

## Abstract

Combining multiple Parkinson’s disease (PD) relevant cellular phenotypes might increase the accuracy of midbrain dopaminergic (mDA) *in vitro* models. We differentiated patient-derived induced pluripotent stem cells (iPSCs) with a LRRK2 G2019S mutation, isogenic control and genetically unrelated iPSCs into mDA neurons. Using automated fluorescence microscopy in 384-well plate format, we identified elevated levels of α-synuclein and Serine 129 phosphorylation (pS129), reduced dendritic complexity, and mitochondrial dysfunction. Next, we measured additional image-based phenotypes and used machine learning (ML) to accurately classify mDA neurons according to their genotype. Additionally, we show that chemical compound treatments, targeting LRRK2 kinase activity or α-synuclein levels, are detectable when using ML classification based on multiple image-based phenotypes. We validated our approach using a second isogenic patient derived SNCA gene triplication mDA neuronal model. This phenotyping and classification strategy improves the exploitability of mDA neurons for disease modelling and the identification of novel PD drug targets.

## Introduction

Parkinson’s disease (PD) is a heterogeneous movement disorder with a combination of motor and non-motor features caused by environmental and genetic risk factors or mutations in specific genes. Pathological characteristics of PD include the progressive loss of midbrain dopaminergic (mDA) neurons and often the appearance of Lewy bodies, cytoplasmic inclusions containing aggregated α-synuclein protein (Blesa et al., 2022; Domingo and Klein, 2018; Poewe et al., 2017).

Mutations in the leucine-rich repeat kinase 2 gene (*LRRK2*) have been associated with PD. The Glycine to Serine substitution at position 2019 (G2019S) in the LRRK2 kinase domain is the most frequent mutation and accounts for 5-6% of familial PD and 1-2% of sporadic cases (Correia Guedes et al., 2010). Increased LRRK2 G2019S kinase activity is believed to be one reason for mDA neuron loss, but the exact mechanism remains unclear (Smith et al., 2006; West et al., 2005; Weykopf et al., 2019). One hypothesis is that LRRK2 G2019S causes defects in mitochondrial biology. Increased autophagy markers, but also PINK1/Parkin-, and Miro1-related defects support the idea that specifically mitophagy-linked processes are disturbed in LRRK2 G2019S neurons (Bonello et al., 2019; Hsieh et al., 2016; Schwab et al., 2017). Additionally, LRRK2 G2019S could induce mDA neuron loss by increasing the levels of phosphorylated α-synuclein, leading to its aggregation, since LRRK2 kinase inhibition can prevent phosphorylated α-synuclein from forming protein inclusions (Daher et al., 2014; Longo et al., 2017; Obergasteiger et al., 2020; Volpicelli-Daley et al., 2016; Xiong et al., 2017).

An emerging picture of PD is therefore that multiple disease mechanisms, such as α-synuclein aggregation or mitochondrial dysfunction act together or can even exacerbate each other. Human patient induced pluripotent stem cell (iPSC)-derived mDA neurons expressing LRRK2 G2019S constitute a valuable *in vitro* model to understand PD pathophysiology and to improve therapeutic hypotheses by experimental testing.

Despite the apparent value of neuronal models, important challenges remain: Individual *in vitro* PD pathological features are often subtle or variable when examined across different differentiation batches or genotypes. Furthermore, single isolated PD phenotypes do not capture the multifactorial complexity of PD. Additionally, and despite their physiologically relevance, iPSC neuronal models are rarely used for PD-related drug discovery due to throughput feasibility concerns based on technical complexity as well as genetic variability (Cobb et al., 2018; Elitt et al., 2018; Farkhondeh et al., 2019). The goal of this study was therefore to develop a robust methodology able to detect multiple cellular PD-related pathophysiological phenotypes in a physiologically relevant human mDA neuronal model system. We aimed for sufficient sensitivity to detect phenotypic variations based on genetic, but also chemical compound induced phenotypic changes.

We demonstrate that multiple PD relevant cellular phenotypes can be detected in microscopic images obtained from human patient LRRK2 G2019S iPSC-derived mDA neurons in 384-well plate format. We show that machine learning (ML) can be used to distinctively classify PD mDA neurons from control neurons based on multiple image-derived phenotypes. Finally, we demonstrate that our multi-phenotype classification approach is sensitive enough to detect different small molecules with a PD-relevant mode of action. We applied our method also to a SNCA gene triplication-carrying mDA neuronal model and obtained similar results. Our work outlines a novel and robust strategy for the use of PD patient iPSC-derived mDA neurons for imaging-based disease modelling by computationally combining multiple disease relevant phenotypes.

## Results

### Differentiation of iPSCs to midbrain dopaminergic cultures

iPSC lines from a patient with a confirmed LRRK2 G2019S mutation and a genetically corrected isogenic control line (GS/+ and +/+, respectively) were differentiated to midbrain dopaminergic (mDA) neural cultures. Differentiated cultures expressed neuronal markers (TUBB3 and MAP2) and dopaminergic neuron markers, including tyrosine hydroxylase (TH) in combination with expression of FOXA2, while the glial marker Glial Fibrillary Acidic Protein (GFAP) was only weakly expressed (**Figure S1A-B**). Immunostaining showed similar percentages of TH- and MAP2-expressing isogenic control +/+ and GS/+ neurons indicating comparable differentiation potentials in both genotypes (**Figure S1C**).

### LRRK2 G2019S mDA neurons overexpress α-synuclein and display mitochondrial dysfunction

To detect image-based hallmarks of PD, we designed an immunofluorescence-based workflow in 384-well plate format. In brief, cryopreserved 30-day old mDA neurons were thawed and seeded in 384-well plates, cultured for 7 days, fixed and stained. Automated microscopy and image segmentation was used to extract multiple quantitative image features. First, GS/+ and +/+ mDA neurons were stained with antibodies against α-synuclein, TH and MAP2. In the TH-positive GS/+ neuronal population, α-synuclein levels were increased by 15% (**Figure 1A**). Western blotting with a different antibody confirmed the increase in α-synuclein levels across multiple differentiation batches (**Figure S2A-D**). In addition, MAP2 staining indicated that less dendritic branches were present in GS/+ neurons (**Figure 1A**). Staining with a pS129 α-synuclein antibody showed that the surface area occupied by pS129 α-synuclein and its fluorescence intensity were increased (**Figure 1B**). To exclude signal originating from non-phosphorylated forms of α-synuclein, we treated the fixed cells with lambda phosphatase. As expected, we observed that lambda phosphatase treatment strongly reduced the pS129 α-synuclein signal intensity in both GS/+ and +/+ neurons (**Figure 1C**). These findings suggest that pS129 α-synuclein levels are indeed increased in GS/+ mDA neurons.

**Figure 1:**
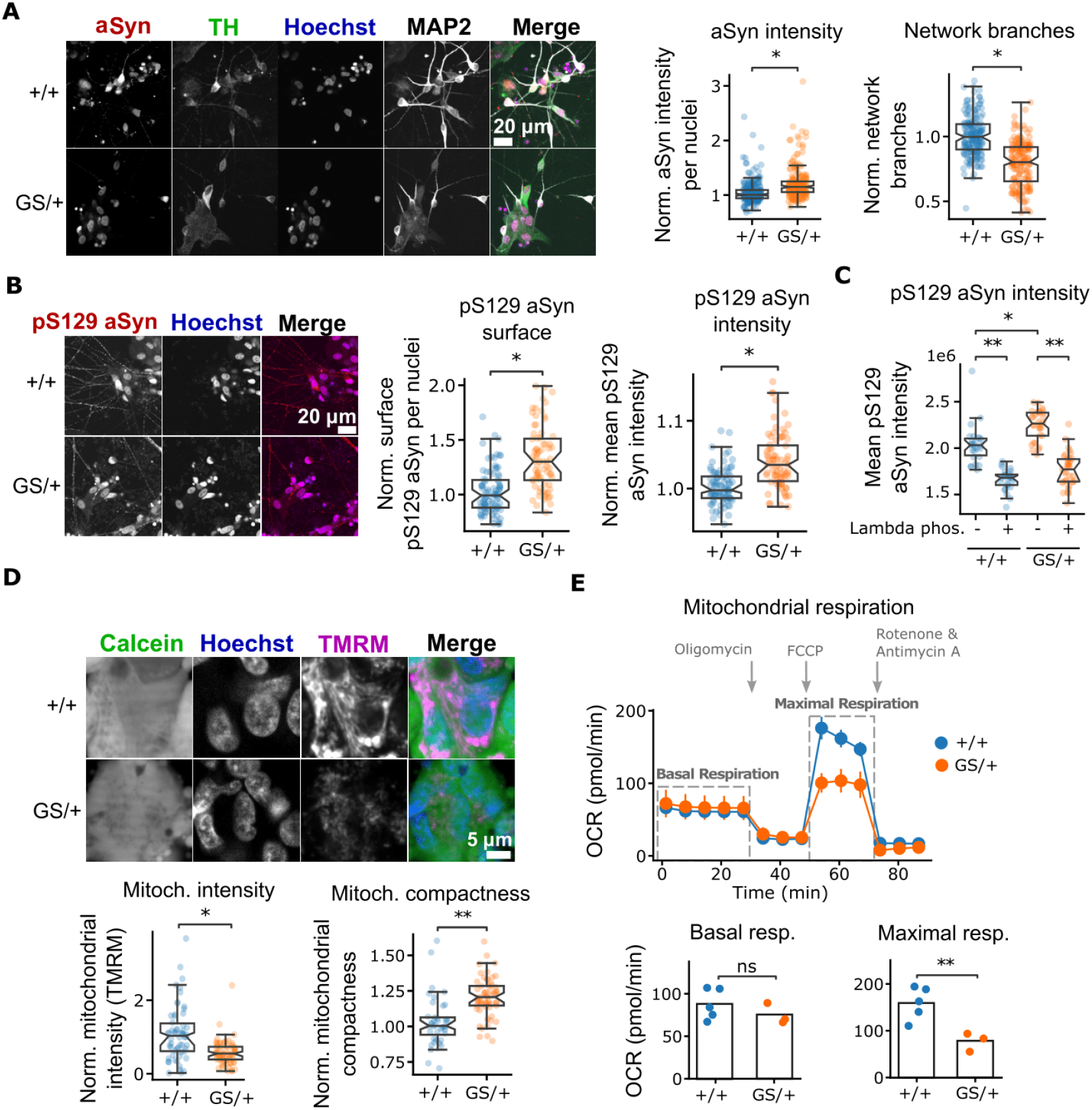
LRRK2 G2019S mDA neurons overexpress α-synuclein and display mitochondrial dysfunction. **(A)** iPSC-derived LRRK2 G2019S mDA neurons were immunostained against α-synuclein, TH and MAP2 and α-synuclein intensity in TH positive GS/+ neurons as well as a neuronal network complexity was quantified in microscopic images. **(B)** Immunofluorescence staining against pS129 α-synuclein, α-synuclein and MAP2 and quantification of pS129 α-synuclein in neurites as well as overall pS129 α-synuclein fluorescence intensity. **(C)** mDANs were treated with lambda phosphatase before staining with a pS129 α-synuclein antibody. **(D)** Staining with the live cell dye Calcein and mitochondrial membrane potential-sensitive dye TMRM and quantification of TMRM intensity and mitochondrial (TMRM) compactness. **(E)** Assessment of mitochondrial respiration using the Seahorse XF analyzer. Seahorse experiments were performed in triplicate and means ± SEMs are shown. Imaging experiments shown in panels **(B)-(D)** were performed at least in duplicate with multiple technical replicates. Each data point represents one well. All data has been median normalized to the respective +/+ condition per plate. Welch’s unequal variances t-test was used for significance testing. Notches in boxplots indicate the 95% confidence interval.

Next, we examined mitochondrial phenotypes. Staining with the live cell dye Calcein and the mitochondrial membrane potential-sensitive dye Tetramethylrhodamine (TMRM) indicated that the overall TMRM fluorescence in living cells was decreased by 33% in GS/+ mDA neurons suggesting that the intactness of the mitochondrial membrane is compromised in GS/+ mDA neurons (**Figure 1D**). Additionally, mitochondria in GS/+ mDA neurons were more compact, indicating an altered morphology compared to the web-like mitochondrial structure in control neurons (**Figure 1D**). We used the mitochondria-targeting toxin Rotenone to chemically validate the TMRM staining (**Figure S3**). Additionally, we measured the oxygen consumption rate (OCR) in our mDA neurons. We found that the basal respiration rate did not differ between both genotypes, while the maximal respiration rate after Carbonyl cyanide p-(tri-fluromethoxy)phenyl-hydrazone (FCCP) treatment was two-fold increased in +/+ control mDA neurons (**Figure 1E**). Taken together, we found that LRRK2 G2019S mDA neurons show multiple hallmarks of PD, including elevated α-synuclein level, presence of pS129 α-synuclein, and mitochondrial dysfunction.

### ML strategy to classify neurons based on image-derived cellular features

We hypothesized that the combination of multiple image-based phenotypes would give rise to a “neuronal fingerprint” and allow the accurate and robust identification of different cell lines or treatment conditions, thereby making it a useful tool for iPSC-based disease modelling or compound screening. We applied different ML algorithms termed “classifiers” to achieve this task. Specifically, we used Linear Discriminant Analysis (LDA) (Fisher, 1936), Support Vector Machine (SVM) (Cortes and Vapnik, 1995), and Light Gradient Boosting Machine (LightGBM) (Ke et al., 2017) algorithms (**Figure 2)**. Quantitative image-derived features were used as input data and the ML classifiers were trained to separate two classes from each other (**Table S1 & 2**). Next, new data (i.e. data from drug treated neurons), was then mapped based on feature similarity to the pre-trained reference classes. For example, to estimate the effect of a chemical compound treatment, compound-treated cells can be classified in comparison to DMSO-treated mutant and wild-type cells. An increased classification proximity generally indicates a higher phenotypic similarity (**Figure 2C**).

**Figure 2:**
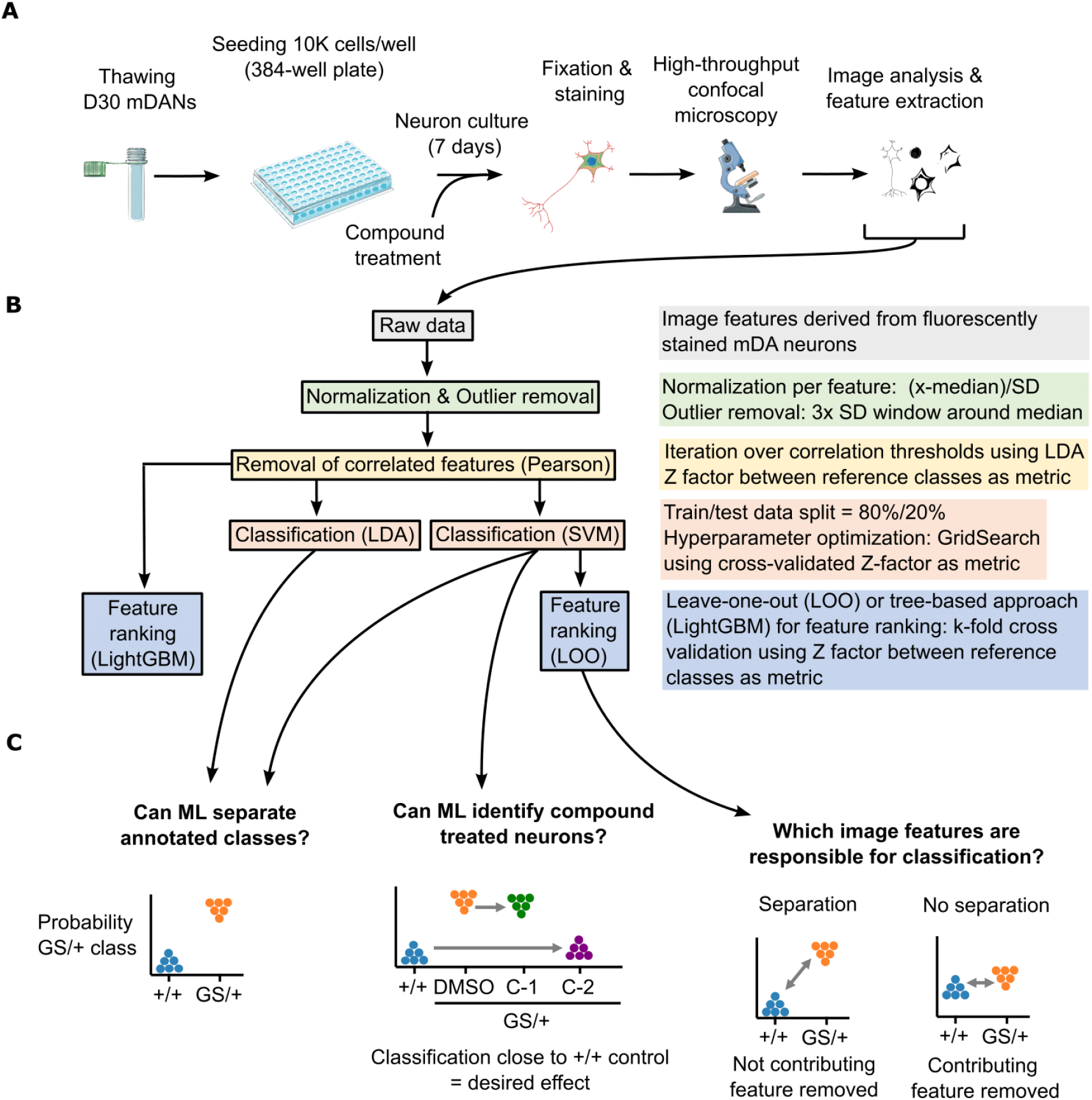
Machine learning (ML) strategy to classify neurons based on image-derived cellular features. **(A)** Schematic depiction of the generation of image-derived cellular feature data. **(B)** Overview of the data processing steps and ML methodology. **(C)** Schematic depiction of how ML classification was used to separate different neuronal cell lines (left panel), identify bioactive chemical compounds (middle panel) and how “leave-one-out” analysis can identify the contribution of individual image-derived cellular features to ML classification (right panel). Whitney U-testing was performed for significance testing.

### ML classification can distinguish neuronal genotypes based on image-derived cellular features

After having identified multiple individual PD-linked phenotypes we asked whether combining a large number of cellular phenotypes would result in a distinct GS/+ phenotypic fingerprint. To evaluate the specificity of a GS/+ phenotypic fingerprint compared to the +/+ isogenic control line, we differentiated two additional and genetically unrelated iPSC lines (EDi001-A-5 and GIBCO) into mDA neurons (**Figure S8**). All four mDA neuronal cell lines were then stained with Hoechst and antibodies against α-synuclein, TH and MAP2. After image acquisition and segmentation, we derived a total of 126 quantitative features from the images (**Table S1**). We hypothesized that a weighted combination of all 126 cellular image features might allow the generation of a unique phenotypic fingerprint per cell line. Secondly, we hypothesized that the generated phenotypic fingerprint of GS/+ neurons would be significantly different from all control mDA neuron control lines.

To test our hypotheses, we evaluated the two supervised ML classifiers LDA and SVM. In a first step, we determined the Pearson correlations of all image features to remove strongly correlated image features (**Figure S4A-C**). Both LDA and SVM algorithms were then trained repeatedly on shuffled sets of 80% of the imaging data and tested on 20% of the imaging data. In total training and testing were repeated 25 times on shuffled slices of the dataset in a process referred to as cross-validation (CV) (**Figure 2B**, **Figure S4D-E**). CV is useful to detect and prevent overfitting and to increase robustness since the ML models are trained on multiple slightly different datasets. We observed that training variability over all cycles was generally low, indicating that sufficient training data was provided to both the LDA and SVM algorithms. Although by eye the four mDA neuronal lines appeared relatively similar (**Figure 3A**), both LDA and SVM classification algorithms successfully distinguished GS/+ neurons from the other three tested cell lines based on the extracted image features. Overall, SVM performed better as indicated by the larger distance between GS/+ neurons and the other three control cell lines (**Figure 3B**). Since GS/+ and its isogenic counterpart +/+ were the two reference classes in our experiment, we calculated the statistical Z-factor between both (Zhang et al., 1999). The SVM classification Z-factor was superior to the LDA Z-factor (0.12 vs. 0.43) (**Figure 3B**). Based on these results, we focused mainly on SVM classification. To obtain a biological meaningful explanation of the classification results, we applied Leave-One-Out Cross-Validation (LOOCV). During LOOCV, each image feature is left out once, classification is performed repeatedly on the remaining image features, and the resulting Z-factor is calculated (**Figure 2C**, *right panel*). LOOCV demonstrated that our SVM results can most likely be explained by cell line differences concerning the ratio of MAP2 positive neurons and the level of α-synuclein (**Figure 3C**). We confirmed the contribution of these image features identified by LOOCV by using LightGBM, a different classification algorithm (**Figure S4F**). Together these findings demonstrate that iPSC-derived mDA neurons can be stably classified by SVM, based on multiple subtle, but detectable, image-extracted phenotypes.

**Figure 3:**
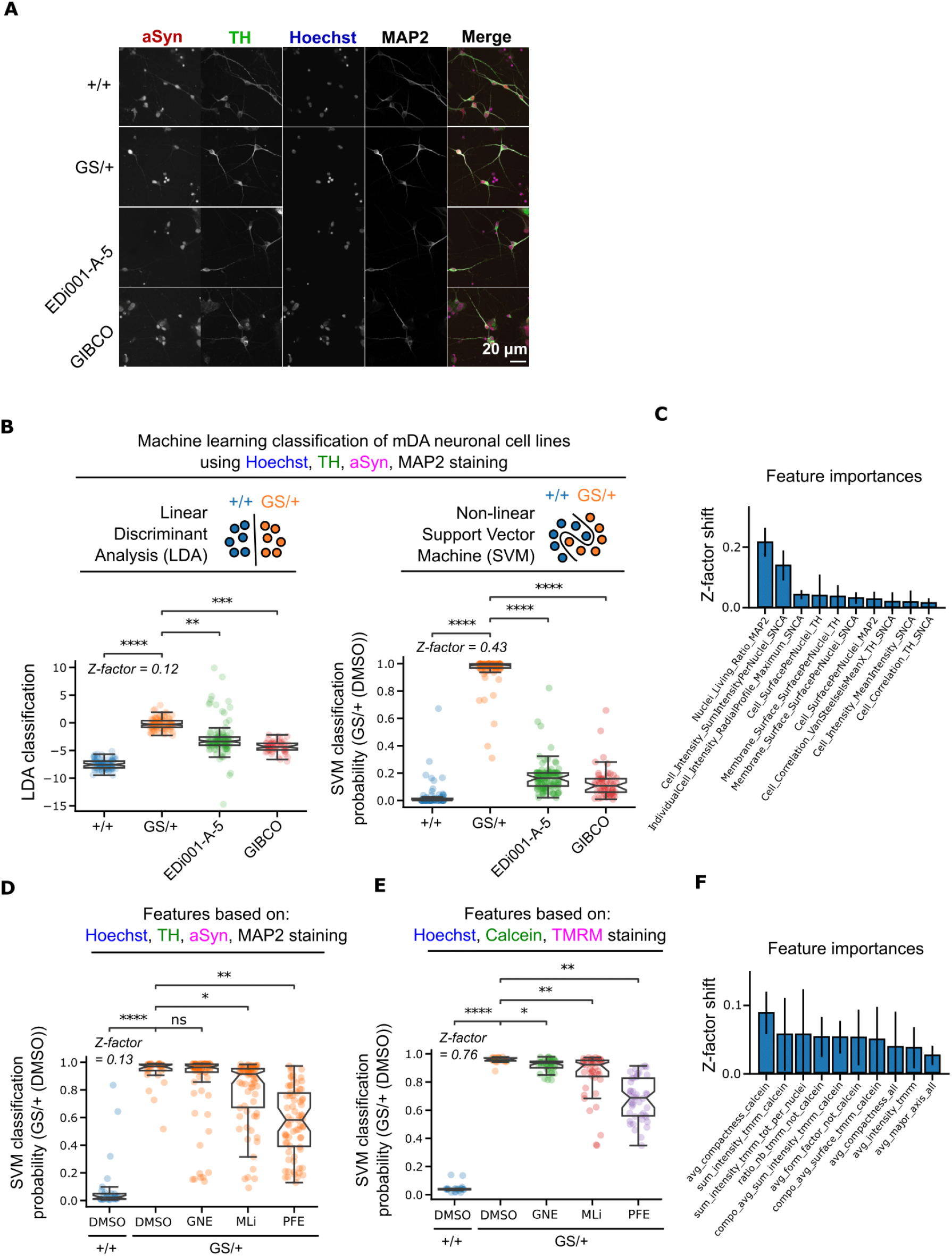
Machine learning (ML) classification can identify genotype-related and chemical compound-induced phenotypic differences based on image-derived cellular features in mDA neurons. **(A)** Representative images of neurons stained with Hoechst and antibodies against TH, α-synuclein and MAP2. Image-derived cellular features were extracted from such images. **(B)** The two supervised ML classification algorithms Linear Discriminant Analysis (LDA) and Support Vector Machine (SVM) were trained to separate the two reference classes GS/+ and +/+ isogenic control mDANs. The additional mDAN control lines were then mapped to the reference classes’ feature space. **(C)** Leave-One-Out Cross-Validation (LOOCV) to identify individual feature contributions to SVM classification of multiple cell lines in **(B)**. **(D)** SVM classification of GS/+ and +/+ isogenic control mDANs and mapping neurons treated with the LRRK2 inhibitors GNE-7915, MLi-2 and PFE-360 to the reference classes’ feature space. **(E)** Same experiment as in **(D)** but instead neurons were stained with Hoechst, Tetramethylrhodamine (TMRM) and Calcein. **(F)** LOOCV to identify individual feature contributions to SVM classification in **(E)**. All imaging data was generated in duplicate experiments with multiple technical replicates. Each data point represents one well. Mann-Whitney U-testing was performed for significance testing. Notches in boxplots indicate the 95% confidence interval.

### Machine learning classification can identify LRRK2 inhibitor treated neurons based on image-derived cellular features

Next, we asked whether SVM-driven analysis is sensitive enough to detect chemical compound induced phenotypic changes. We hypothesized that LRRK2 inhibitor treatment might partially rescue the previously observed combined feature phenotype (**Figure 3B**). Cryopreserved D30 mDA neurons were thawed and seeded in 384-well plates. Five days after seeding the LRRK2 inhibitors GNE-7915, PFE-360, and MLi-2 were added for 48 hours and the neurons were fixed and stained using Hoechst, α-synuclein, TH, and MAP2 antibodies. Image-based feature extraction, data processing and SVM model training were performed (**Figure S5**). The SVM classifier successfully distinguished +/+ and GS/+ mDA neurons treated with DMSO with a Z-factor of 0.13 (**Figure 3D**). Next, LRRK2 inhibitor treated GS/+ mDA neurons were classified relative to the DMSO controls. GNE-7915 did not lead to phenotypic changes detectable in our assays and resembled the DMSO control classification. PFE-360 and MLi-2 induced subtle phenotypic differences detected by Hoechst/α-synuclein/TH/MAP2 staining and were classified as significantly different from DMSO-treated neurons. The shift towards the +/+ isogenic control was strongest for the PFE-360 treated GS/+ mDA neurons (**Figure 3D**).

Next, we tested whether LRRK2 inhibitor treatment would also lead to SVM-detectable multiphenotypic changes on the mitochondrial level. Neurons were cultured and treated as before and stained with Hoechst, the live cell dye Calcein and the mitochondria-specific dye TMRM. We extracted a total of 96 mitochondria-related image features based on these three stainings (**Table S2**) and trained a SVM model using these features to distinguish +/+ from GS/+ mDA neurons (**Figure S6**). We then applied the SVM model to sets of mitochondrial image features from LRRK2 inhibitor-treated mDA neurons. Similar to the previous results obtained with the Hoechst/α-synuclein/TH/MAP2 staining, we detected only a weak effect of GNE-7915 on the measured mitochondrial phenotypes, while PFE-360 and MLi-2 treatment of GS/+ mDA neurons led to a classification shift towards +/+ isogenic control mDA neurons (**Figure 3E**). To identify the mitochondrial features most responsible for the observed classification result, we performed LOOCV analysis. We found that mitochondrial shape (i.e. compactness and form factor) as well as TMRM intensity contributed the most to the classification result (**Figure 3F**).

### Detection PKC agonist-treated single wells using multiple image-derived cellular features in LRRK2 G2019S neurons

Recently, Laperle *et al*. demonstrated that lysosomal activation by phorbol esters, such as PEP005 and Prostratin, reduced α-synuclein levels in iPSC-derived mDA neurons (Laperle et al., 2020). Given the established connection between LRRK2 and lysosomal biology, we hypothesized that PEP005 and Prostratin might also be able to lower the elevated α-synuclein levels in our LRRK2 G2019S model and thereby shift multiple cellular phenotypes towards an unmutated control phenotype (Hockey et al., 2015; Obergasteiger et al., 2020). To demonstrate that ML classification is sensitive enough to detect a chemical modulation in mDA neurons, we tested the sensitivity of our model by treating only 6 randomly selected wells per biological replicate with PEP005 or Prostratin for 72 hours (**Figure 4A**). Next, cells were fixed and stained with Hoechst and α-synuclein, TH and MAP2 antibodies. After microscopic imaging and segmentation, 126 image features were extracted (**Table S1**). Verification of individual image features, such as the number of TH-positive cells, showed that PEP005 and Prostratin compound treatment was neither toxic for GS/+ nor +/+ neurons (**Figure 4B**). Confirming our previous results, we detected elevated α-synuclein levels in DMSO-treated GS/+ neurons in this experimental setup. Importantly, PEP005 and Prostratin treatment led to a statistically significant decrease in α-synuclein levels specifically in GS/+ neurons, but not control +/+ neurons, confirming the initial results of Laperle *et al*. obtained in different PD mDA neuron lines (**Figure 4C**). Next, we trained a SVM model to distinguish +/+ from GS/+ mDA neurons using image-based features as input (**Figure S7**). Consistent with our previous results, SVM was able to separate both DMSO-treated control classes +/+ and GS/+ with high accuracy (0.98 ± SEM 0.02) and a Z-factor of 0.72 (**Figure 4D**). We then applied the SVM model to sets of image-features originating from PEP005 and Prostratin treated wells. PEP005 and Prostratin treated GS/+ neurons classified differently than the DMSO-treated GS/+ neurons. Although this effect was small for PEP005, most Prostratin treated wells shifted towards the +/+ isogenic control neurons. Additionally, we observed that compound treated +/+ control neurons responded less to PEP005 and Prostratin treatment (**Figure 4D**).

**Figure 4:**
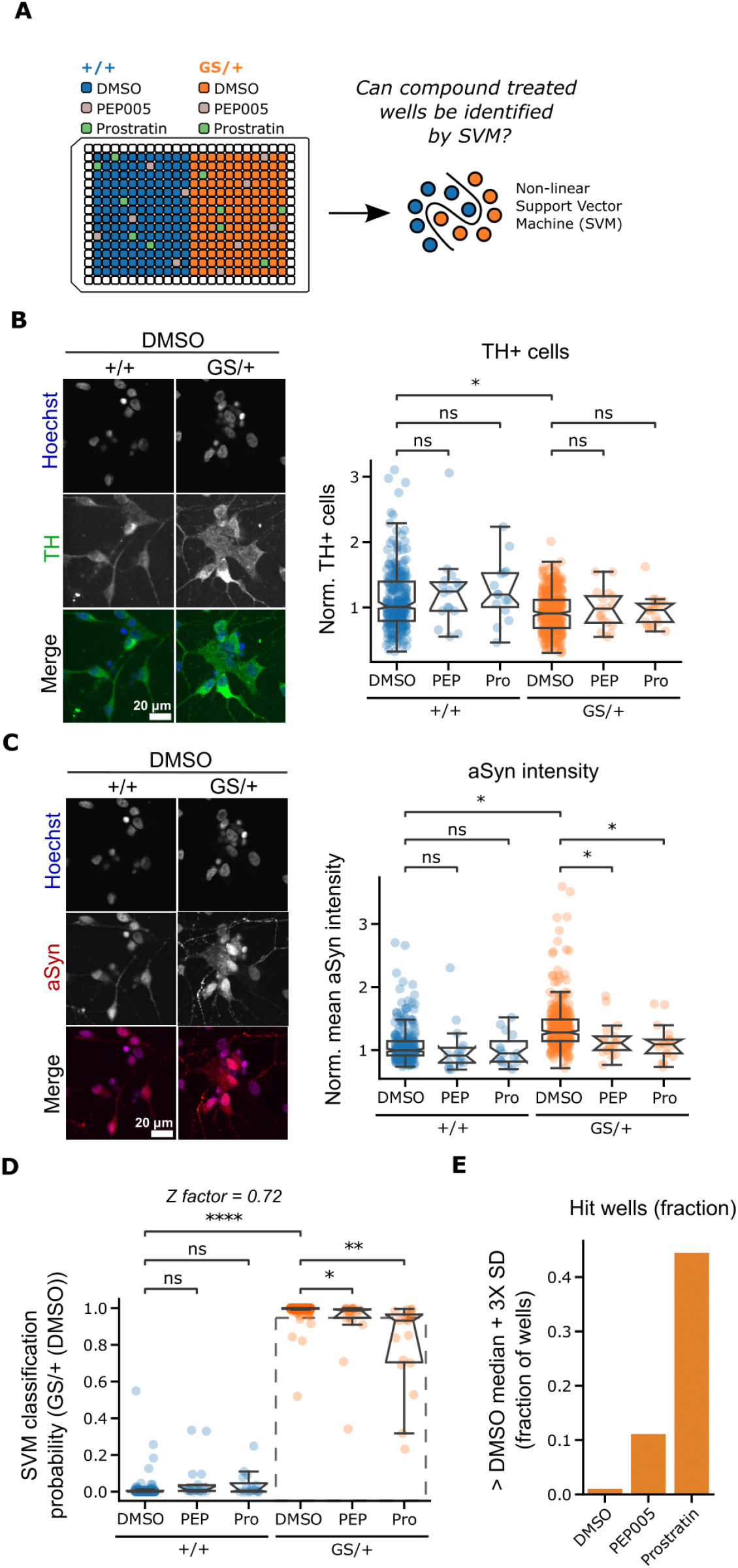
Machine learning (ML) can identify PKC agonist treated LRRK2 G2019S mDA neurons in a simulated screening setup. **(A)** Schematic depiction of experimental design. Single wells spiked with PEP005 or Prostratin were randomly distributed over the plate. Support Vector Machine (SVM) classification was applied to identify these wells. **(B)** and **(C)** Representative images illustrate PEP005 and Prostratin effects on the number of Tyrosine hydroxylase (TH) positive cells and α-synuclein staining intensity. **(D)** SVM classification of GS/+ and +/+ isogenic control mDANs based on cellular image features extracted from Hoechst, α-synuclein, TH, and MAP2 staining. PEP005 and Prostratin treated wells were then mapped to the references classes’ feature space. The broken square includes datapoints (wells) that are more than three standard deviations (SD) from the GS/+ DMSO treated median. **(E)** Quantification of the fraction wells more than three SDs from the GS/+ DMSO treated class median. All imaging data was generated in triplicate experiments with multiple technical replicates. Each data point represents one well. Mann-Whitney U-testing was performed for significance testing. Notches in boxplots indicate the 95% confidence interval.

To assess whether single PEP005 or Prostratin treated wells could be detected in a typical screen setup using only a small number of replicates, we determined a 3x SD threshold around the median of the DMSO-treated GS/+ neurons. Next, we calculated the percentage of compound treated wells beyond the threshold that could be regarded as a “hit”. For GS/+ neurons treated with DMSO less than 1% of wells were more than 3 SDs away from the median, while this was 11% of PEP005 and 43% of Prostratin treated wells (**Figure 4E**).

### Detection PKC agonist-treated single wells using multiple image-derived cellular features in SNCA triplication neurons

To generalize our multi phenotype approach, we established a second PD mDA neuron model based on SNCA gene triplication-carrying donor iPSCs expressing four copies of SNCA. Additionally, we differentiated isogenic control iPSCs expressing two SNCA copies into mDA neurons (**Figure S8**). Using both cell lines, we performed a similar experiment as described in **Figure 4A** with the aim to detect individual wells treated with PEP005 or Prostratin using SVM classification (**Figure 5A**). SNCA triplication mDA neurons showed signs of α-synuclein accumulation in dendrites and a reduced dendritic network (**Figure 5B**). Image feature quantification confirmed, that indeed α-synuclein levels were increased in SNCA triplication mDA neurons. Additionally, we observed α-synuclein lowering effects of 15% by PEP005 and 25% by Prostratin (**Figure 5C**). Next, we trained a SVM classifier to separate isogenic control from SNCA triplication mDA neurons. Like our previous findings using the LRRK2 model, the SVM algorithm was able to separate isogenic control from SNCA triplication mDA neurons with high accuracy (0.97 ± SEM 0.03) resulting in a Z-factor of 0.73 (**Figure 5D**, **Figure S9**). SVM classification of SNCA triplication mDA neurons treated with PEP005 or Prostratin showed a shift towards isogenic control mDA neurons. Isogenic control neurons treated with both compounds had a similar image feature-based profile and were statistically indistinguishable from DMSO-treated control neurons, suggesting a specific effect of PEP005 and Prostratin in SNCA triplication neurons (**Figure 5D**).

**Figure 5:**
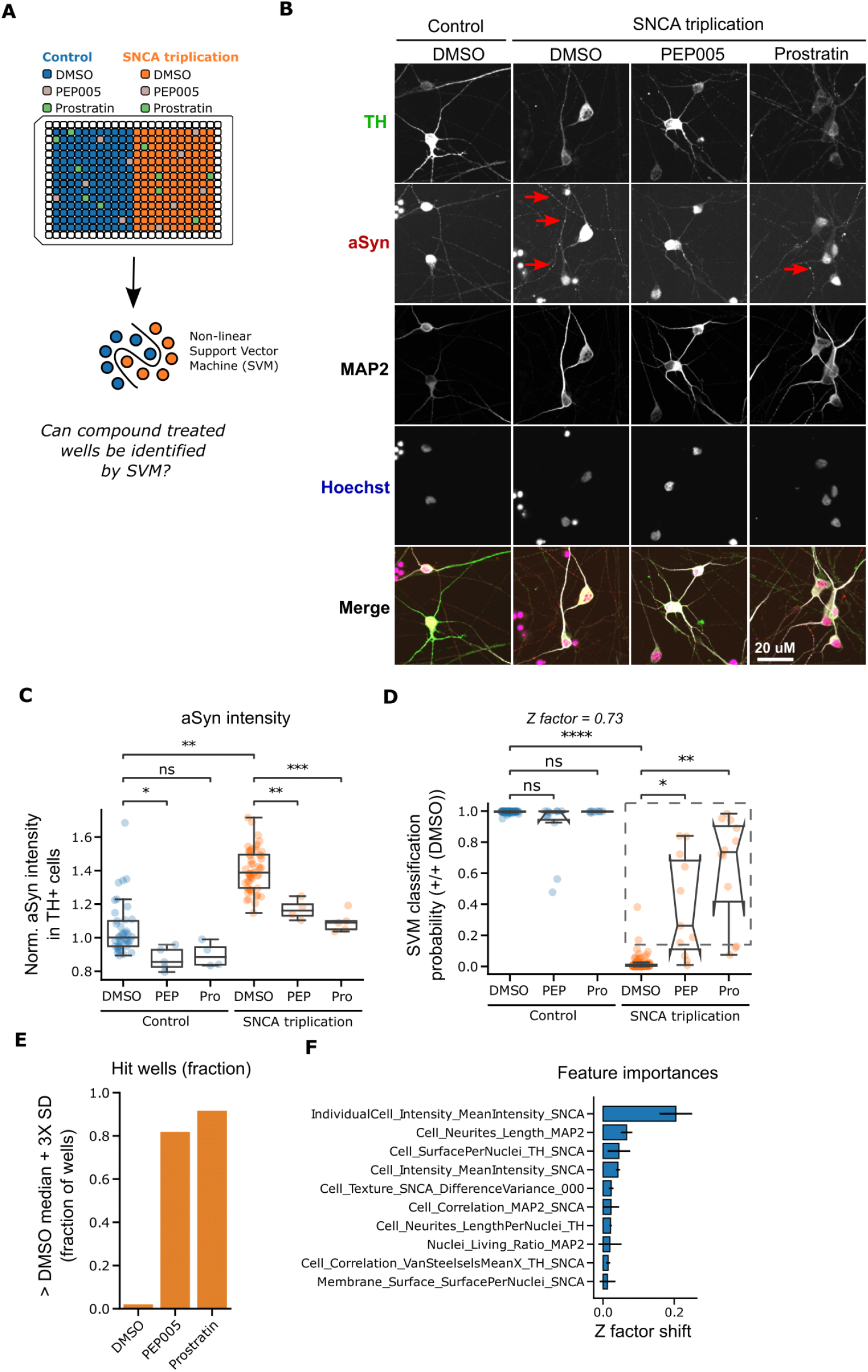
Machine learning (ML) can identify Protein Kinase C (PKC) agonist treated SNCA triplication mDA neurons in a simulated screening setup. **(A)** Schematic depiction of experimental design. Single wells spiked with PEP005 or Prostratin were randomly distributed over the plate. Support Vector Machine (SVM) classification was applied to identify these wells. **(B)** Representative images of neurons stained with Hoechst, and TH, α-synuclein and MAP2 antibodies after 37 days of differentiation and treated with either DMSO, PEP005 or Prostratin. Red arrows indicate α-synuclein staining in neurites. **(C)** Quantification of α-synuclein staining intensity across all treatment conditions. **(D)** SVM classification of SNCA triplication and isogenic control mDANs based on cellular image features extracted from Hoechst, α-synuclein, TH, and MAP2 staining. PEP005 and Prostratin treated wells were then mapped to the reference classes’ feature space. The broken square includes datapoints (wells) that are more than three standard deviations (SD) from the SNCA triplication DMSO treated median. **(E)** Quantification of the fraction of wells more than three SDs from the SNCA triplication DMSO treated class median. **(F)** Leave-One-Out Cross-Validation (LOOCV) to identify individual feature contributions to SVM classification in **(D)**. All imaging data was generated in triplicate experiments with multiple technical replicates. Each data point represents one well. Mann-Whitney U-testing was performed for significance testing. Notches in boxplots indicate the 95% confidence interval.

To assess whether single PEP005 or Prostratin treated wells could be detected in a typical screen setup using only a small number of replicates, we again determined a 3x SD threshold around the median of the DMSO-treated SNCA triplication neurons and calculated the percentage of compound treated wells beyond the threshold. For SNCA triplication neurons treated with DMSO less than 1% of wells were more than 3 SDs away from the median, while this was 81% of PEP005 and 91% of Prostratin treated wells (**Figure 5E**). To deduce which image features contributed to the successful separation of SNCA triplication and isogenic control mDA neurons by SVM classification we performed LOOCV. As expected, the single most important image feature distinguishing both cell lines was the α-synuclein staining intensity, a proxy for cellular α-synuclein content (**Figure 5F**). This feature explained 0.2 points of the observed 0.73 Z-factor.

We confirmed the contribution of α-synuclein content and other features by using LightGBM, a different classification algorithm (**Figure S9F**). These findings in a second PD-relevant disease model indicate that bioactive molecules such as PEP005 and Prostratin can be detected using our multiphenotypic approach and a relatively small number of technical replicates.

## Discussion

In this study, we demonstrate that image-derived phenotypes in human iPSC-derived mDA neurons can be used for cell line stratification and the identification of chemical compound treated neurons by ML classification approaches. iPSC-derived neurons are only rarely used in drug discovery due to complex cell culture protocols, long culture duration, and genetic or clonal heterogeneity (Cobb et al., 2018; Elitt et al., 2018; Farkhondeh et al., 2019). We applied multiple strategies to improve the reproducibility of our iPSC-derived neuron models. First, we worked with large cryopreserved batches in order to reduce the number of required differentiations. We also used LRRK2 G2019S and SNCA triplication mDA neurons with their respective isogenic controls in order to reduce sources of inter-donor genetic variability. Additionally, we developed a compact seven day experimental protocol in 384-well plate format to reduce intervention steps related to cell culturing or compound treatment. To further minimize sources of technical variability we semi-automated key cell handling steps and imaging.

The functions of LRRK2 are not fully understood, but it has become clear that LRRK2 can trigger autophosphorylation at Ser1292 and phosphorylate a subset of Rab small GTPases (Rab8A and Rab10) (Rocha et al., 2022; Sheng et al., 2012; Steger et al., 2016). A direct readout of these targets was not present in our panel of stains. This is likely the reason why one of the three tested LRRK2 inhibitors showed only little effects in our experimental setup. Similarly, the used phorbol esters PEP005 and Prostratin have specific phosphorylation inducing effects on PKC subunits α and δ (Hampson et al., 2005; Laperle et al., 2020; Mischak et al., 1993), which we did not examine directly in our phenotypic characterization. Despite this, we observed PEP005 and Prostratin effects in both the LRRK2, but especially the SNCA triplication model, likely because both molecules have α-synuclein lowering capabilities in mDA neurons (Laperle et al., 2020). Since we wanted to capture a broad panel of PD-relevant phenotypes to remain target agnostic with respect to novel modes of actions, we purposely did not include LRRK2 or PKC specific readouts into our cellular staining protocols to reduce target bias.

A small number of studies describe small molecule screening in iPSC-derived neurons in the context of neurodegenerative disorders (Imamura et al., 2017; Kondo et al., 2017; Tabata et al., 2018; Yamaguchi et al., 2020). Of those, Tabata *et al*. and Yamaguchi *et al*. used iPSC-derived DA neurons. Tabata *et al*. screened 1165 FDA-approved drugs and used resistance to Rotenone-induced apoptosis and neurite outgrowth as phenotypic readouts (Tabata et al., 2018). Yamaguchi *et al*. screened 320 compounds and used resistance to carbonyl cyanide m-chlorophenylhydrazone (CCCP)-induced apoptosis and rescued mitophagy as readouts (Yamaguchi et al., 2020). We also observed hallmarks of PD in our mDA neurons such as increased α-synuclein levels, S129 phosphorylation and mitochondrial dysfunction. In contrast to previous work, we use ML classification to bundle multiple phenotypes which offers certain advantages: The used cellular stainings allow the extraction of a large number of PD-relevant image features and thereby create a more biologically diverse representation of mDA neurons amendable to chemical interventions. Second, the combination of multiple, including subtle, phenotypes is statistically more robust than single phenotypic approaches. Additionally, our ML classification approach allows to determine which phenotypic features contributed in particular to the overall phenotypic differences between healthy and disease mDA neurons and might therefore aid the target deconvolution process.

In summary, we developed an experimental and analytical framework using image-based multidimensional readouts capturing multiple PD relevant phenotypes in mDA neurons. We anticipate that this approach could increase the chance to detect active chemical compounds which rescue not only an isolated phenotype, but an ensemble of disease relevant phenotypes.

## Experimental procedures

### Generation of iPSC lines & differentiation into mDA neurons

All iPSC lines were generated by third parties and are deposited in the European Bank for Induced Pluripotent Stem Cells (EBiSC, https://cells.ebisc.org/) and listed in the Human Pluripotent Stem Cell Registry (hPSCreg, https://hpscreg.eu/) (**Table S3**). The original generators have obtained the informed consent from the donors. iPSCs were cultivated on Geltrex-coated (Thermo Fisher Scientific) dishes in StemMACS iPS-Brew XF (Miltenyi Biotech). The medium was changed daily, and cells were passaged twice a week using 0.5 mM EDTA in PBS (Thermo Fisher Scientific). Mycoplasma testing was performed twice per month.

mDA neurons were differentiated using a modified protocol based on Kriks *et al*. (Kriks et al., 2011; Ryan et al., 2013; Weykopf et al., 2019). Briefly, iPSCs were seeded onto Geltrex-coated 6 well plates or T75 flasks at a density of 2×10^5^ cells/cm2 in StemMACS iPS-Brew XF containing 10 μM Y-27632 (Hiss). The next day, medium was switched to KnockOut DMEM medium containing KnockOut serum replacement (both Thermo Fisher Scientific) supplemented with 200 nM LDN19318 (Axon Medchem) and 10 μM SB431542 (Biozol) for dual SMAD-inhibition. On day 2, also 100 ng/ml Shh C24II (Miltenyi Biotech), 2 μM Purmorphamine (Miltenyi Biotec), 100 ng/ml FGF8 (Peprotech) and 3 μM CHIR99021 (Miltenyi Biotec) were added to the medium. After 5 days, medium was gradually shifted to Neurobasal medium (Thermo Fisher Scientific) and SB431542 was omitted from the medium. Starting at day 7, cells were grown only in the presence of LDN19318 and CHIR99021. On day 11, cells were switched to Neurobasal/B27/L-glutamine medium supplemented with CHIR99021 only. On day 13, cells were replated onto Geltrex-coated dishes in Neurobasal/B27/L-glutamine medium supplemented with 20 ng/ml BDNF, 20 ng/ml GDNF (both Cell Guidance Sys.), 221 μM L-ascorbic-acid (Sigma-Aldrich), 10 μM DAPT (Axon Medchem), 1 ng/ml TGF-ßIII (Peprotech), 0.5 mM dibutyryl-cAMP (Enzo Life Sciences) and 10 μM Y-27632 (Hiss). Cells were maintained in the same medium but without Y-27632. Around day 23 – 25, cells were dissociated using StemPro Accutase (Thermo Fisher Scientific) and plated at a density of 1.4 x 105 cells/cm2 onto Geltrex-coated dishes. To eliminate non-neuronal cells, cultures were treated with 1 μg/ml Mitomycin C for 2 hours on day 26. At day 30, neuronal cultures were dissociated using StemPro Accutase supplemented with 10 μM Y-27632 and singularized. Cells were counted and cryopreserved at 2.5×10^6^ cells/vial in CryoStor CS 10 (Sigma Aldrich).

### Neuronal culture & compound treatment

Cryopreserved 30 DIV (days *in vitro*) old neurons were thawed in a water bath and centrifuged (400g, 5 min, RT) in basal medium (**Table S3**) supplemented with ROCK inhibitor (Miltenyi, #130-095-563). Cell pellets were resuspended in differentiation medium (**Table S3**) supplemented with ROCK inhibitor. 384-well plates (Perkin Elmer, #6007558) were coated with 15 μg/ml Poly-L-Ornithin for 1 hour at 37 °C followed by 10μg/ml Laminin overnight at 4 °C. Using Tryphan Blue (Sigma, # T8154-20ML) and a Countess automated cell counter (Invitrogen) 10×10^3^ cells/well were seeded in 384-well plates. Edge wells were avoided for seeding and filled with PBS. Typically, thawed cells were incubate at 37 °C and 5 % CO_2_ for seven days until 37 DIV with differentiation medium changes every other day. Plate coating, cell seeding and medium changes were initially performed manually by multichannel pipetting and later automated using an Agilent Bravo pipetting robot (Agilent) and EL406 plate washer and dispenser (Biotek). Compound treatment with 1μM PEP005 (Tocris, #4054) and 5μM Prostratin (Tocris, #5749) was performed five days after thawing at 35 DIV for 72 hours until 37 DIV. Compound treatment with 0.1μM GNE-7915 (MedChemExpress, #HY-18163), 0.1μM MLi-2 (MedChemExpress, #HY-100411), and 0.1μM PFE-360 (MedChemExpress, #HY-120085)) was performed six days after thawing at 36 DIV for 48 hours until 37 DIV. Treatment with 0.1μM Rotenone (Sigma, #R8875) was performed for 24 hours until DIV 37. For Western blotting experiments, 5μM AraC (Cytosine β-D-arabinofuranoside hydrochloride, Sigma, #C6645) was added for 24 hours before cell lysis on DIV 37 or 44.

### *In situ* cytochemistry

Fixation was performed in 4% paraformaldehyde (PFA, EMS Euromedex, #15710) for 20 minutes, followed by two PBS (Gibco, #14190) washes and permeabilization and blocking with 10% FBS (Gibco, #10270-106) and 0.1 % Triton X-100 (Sigma, #T9284) dissolved in PBS for 30 minutes. Primary antibodies (**Table S3**) were prepared in antibody dilution buffer (PBS supplemented with 5% FBS and 0.1 % Triton X-100) and incubated with the cells overnight at 4°C, followed by three PBS washes. Secondary antibodies and Hoechst (**Table S3**) in antibody dilution buffer were added to the cells for 1 hour at RT, followed by three PBS washes. Mitochondrial imaging was performed in live cells. All dyes (**Table S3**) were prepared in differentiation medium and incubated with the cells for 30 minutes at 37 °C and 5 % CO_2_, followed by a wash with differentiation medium. Cells were imaged in a preheated microscope chamber at 37°C and 5% CO_2_. *In situ* cytochemistry was initially performed manually by multichannel pipetting and later automated using an Agilent Bravo pipetting robot (Agilent) and EL406 plate washer and dispenser (Biotek).

### Imaging & image analysis

All imaging experiments were performed on a Yokogawa CV7000 microscope in scanning confocal mode using a dual Nipkow disk. 384-well plates (Perkin Elmer, #6007558) were mounted on a motorized stage and images were acquired in a row-wise “zig-zag” fashion at RT for fixed cells and 37°C and 5% CO_2_ for living cells. The system’s CellVoyager software and 405/488/561/640nm solid laser lines were used to acquire single Z-plane 16-bit TIFF images through a dry 40X objective lens using a cooled sCMOS camera with 2560×2160 pixels and a pixel size of 6.5μm without pixel binning. Nine images in a 3×3 orientation were acquired from the center of each well. Image segmentation and feature extraction was performed with an in-house software written in C++. Except for the detection of mitochondrial structures, image segmentation was performed on illumination corrected raw images based on fluorescent channel intensity thresholds empirically determined per plate. Multiple quantitative image features were calculated (**Table S1 & S2**). Mitochondrial structures and features were detected in rolling-ball background subtracted and top-hat filtered images similar to a protocol described previously (Iannetti et al., 2016).

### ML analysis

To support the reproducibility of the ML method of this study, the ML summary table is included in the **Supplemental Information** as per data, optimization, model and evaluation (DOME) recommendations (Walsh et al., 2021) (**Table S4**). Multiple datasets were generated differing in terms of the used mDA neurons, chemical compound treatment, fluorescent staining and extracted image features (**Table 1**). The input data was normalized, outliers were removed, and the number of input features was reduced by removing strongly correlated features (**Figure 2**, **Table S4**). We applied the Python-written ML library scikit-learn to train and test all models (Pedregosa et al., 2011). We used predominantly supervised binary classification algorithms with a focus on the non-linear SVM algorithm. **Figure 2B** summarizes the overall ML workflow. All models’ hyperparameters were systematically optimized using scikit-learn’s GridSearchCV module. All models were evaluated using k-fold cross-validation and performance was checked using accuracy. All raw data can be found in the **Supplemental Information.** ML pipelines are available as Jupyter notebooks on GitHub (https://github.com/johanneswilbertz/mDA-neuron-classification).

**Table 1:**
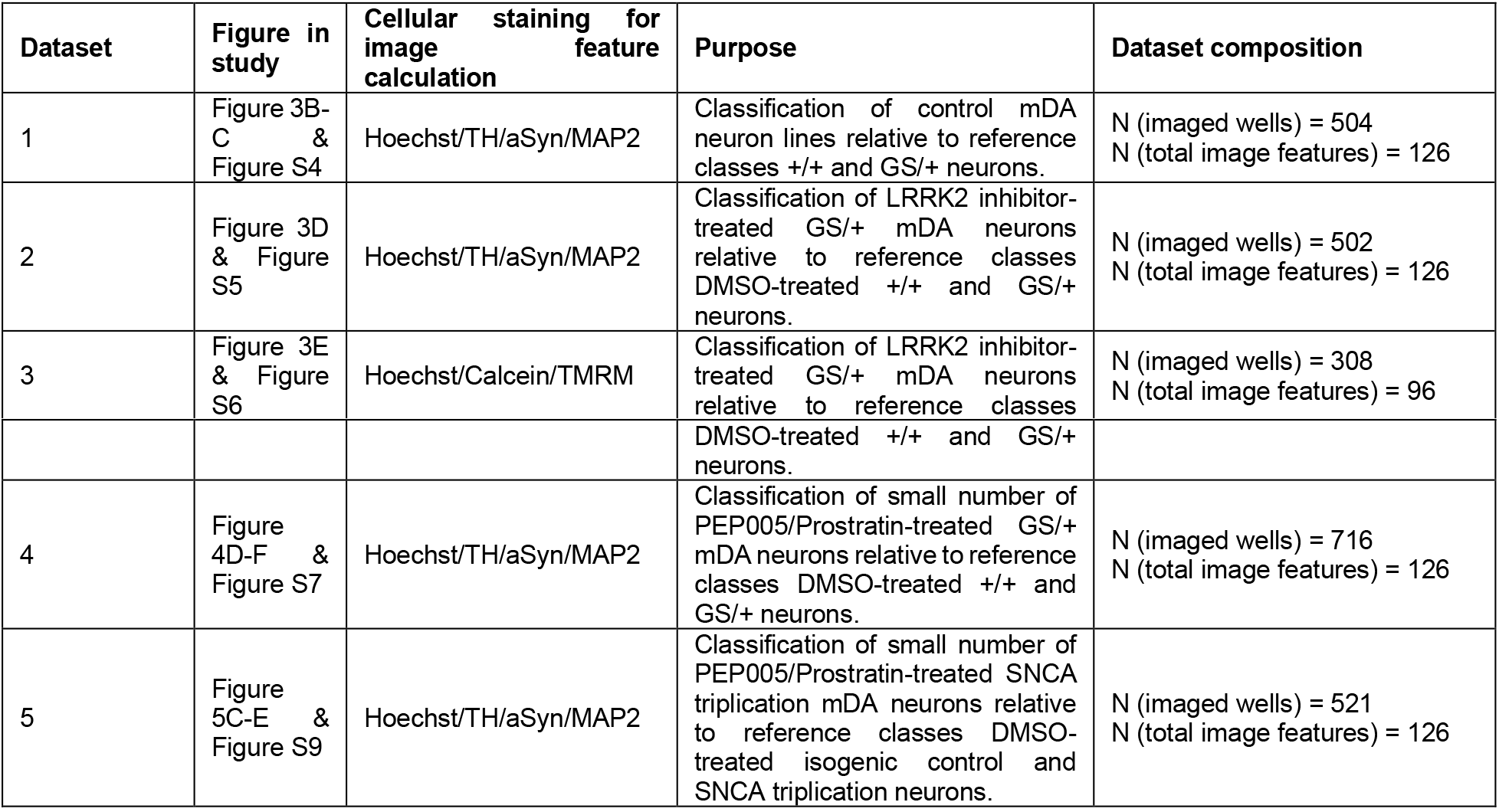
Overview of generated data sets for ML analysis.

### Statistics

All data was generated at least in duplicate with neurons from a single differentiation batch being cultured, stained, and imaged in separate plates and on different days. All data are represented as boxplots including all data points. Each data point represents the mean of a single 384-well plate well comprised of 9 images. Each boxplots’ inner box represent 2^nd^ – 3^rd^ quartile of the data. The horizontal line inside the box represents the median. The notches of box represent the 95% confidence interval of the median obtained by bootstrapping with parameter value 1,000. Boxplot whiskers represent 1.5x of the 2^nd^ – 3^rd^ inner quartile range. Data from different plates was median normalized to allow comparison across plates acquired on different days. Data processing and plotting was carried out with Python packages Pandas (McKinney, 2010), Matplotlib (Hunter, 2007) and Seaborn (Waskom, 2021). Null-hypothesis significance testing was performed with the freely available Python package Statannot (Weber, 2022). For data not displaying a normal distribution, the non-parametrical Mann-Whitney U-test was performed. For normally distributed data, Welch’ t-test was applied. Statistical significance is presented in the figures as * = p < 0.05, ** = p < 0.01, *** = p < 0.001, **** = p < 0.0001, and not significant (ns = p > 0.05).

## Author contributions

AV performed neuronal differentiation, imaging and biochemical experiments; LC developed the image analysis algorithms, performed image analysis and data processing; ZH developed the ML pipeline; NWD developed the image analysis software; BW, MS and SH performed neuronal differentiation and quality controls; MP and OB supervised and designed neuronal differentiation procedures; AO supervised the development of image analysis and data processing algorithms; PS and LB co-supervised the study and designed experiments; JHW co-supervised the study, designed experiments, performed image analysis and data processing, and wrote the manuscript with input from all authors.

## Acknowledgements

The authors thank Bruno Dos Santos (Luxembourg Centre for Systems Biomedicine) for help with Seahorse experiments and the entire Ksilink team for input during project discussions.

## Declaration of interests

The authors declare no competing interests.

## Notes

https://github.com/johanneswilbertz/mDA-neuron-classification

### Competing Interest Statement

The authors have declared no competing interest.

