## Supplemental Data 1 for "Machine learning-aided multidimensional phenotyping of Parkinson’s disease patient stem cell-derived midbrain dopaminergic neurons"

### Supplemental Information

#### Figure S1

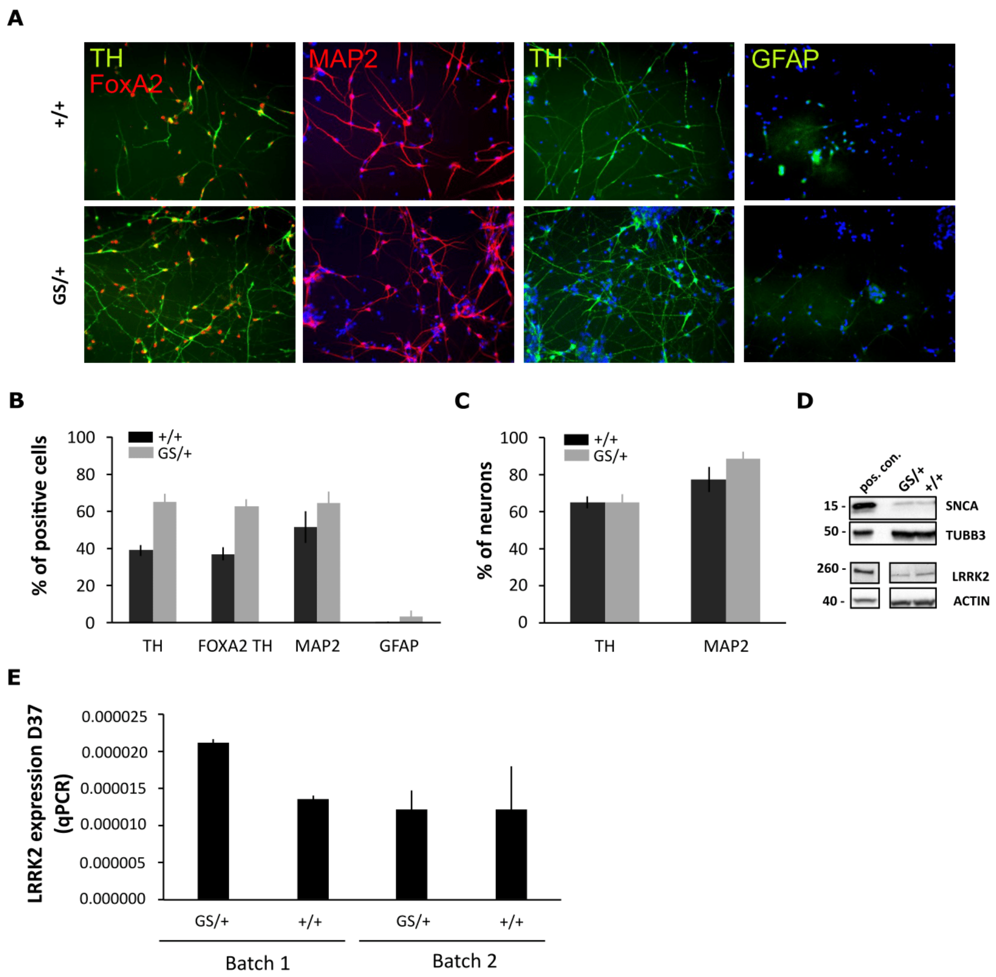


**Figure S1 (referring to Figure 1): Characterization of human iPSC-derived LRRK2 G2019S mDA neurons.** (**A**) Batches of +/+ isogenic control and GS/+ iPSCs were differentiated into mDA neurons for 30 days and cryo-preserved at 2x106 cells per well. Thawed neurons were cultured for additional 7 days and stained for Tyrosine hydroxylase (TH), Forkhead box protein A2 (FOXA2), Microtubule-associated protein 2 (MAP2), and Glial Fibrillary Acidic Protein (GFAP). (**B**) The purity of the mDA neuron population after differentiation and thawing was measured by the ratio of neuronal marker MAP2, dopaminergic marker TH, midbrain dopaminergic markers TH/FOXA2, and astrocyte marker GFAP positive cells. (**C**) The ratio of TH and MAP2 positive cells was calculated for the β-tubulin III (TUBB3) staining-positive neuronal population. (**D**) The presence of the proteins α-synuclein, TUBB3, and LRRK2 in the neuronal culture was confirmed by Western blotting. (**E**) qPCR analysis showed the expression of LRRK2 in isogenic control +/+ neurons as well as in mutation carrying GS/+ neurons.

### Figure S2

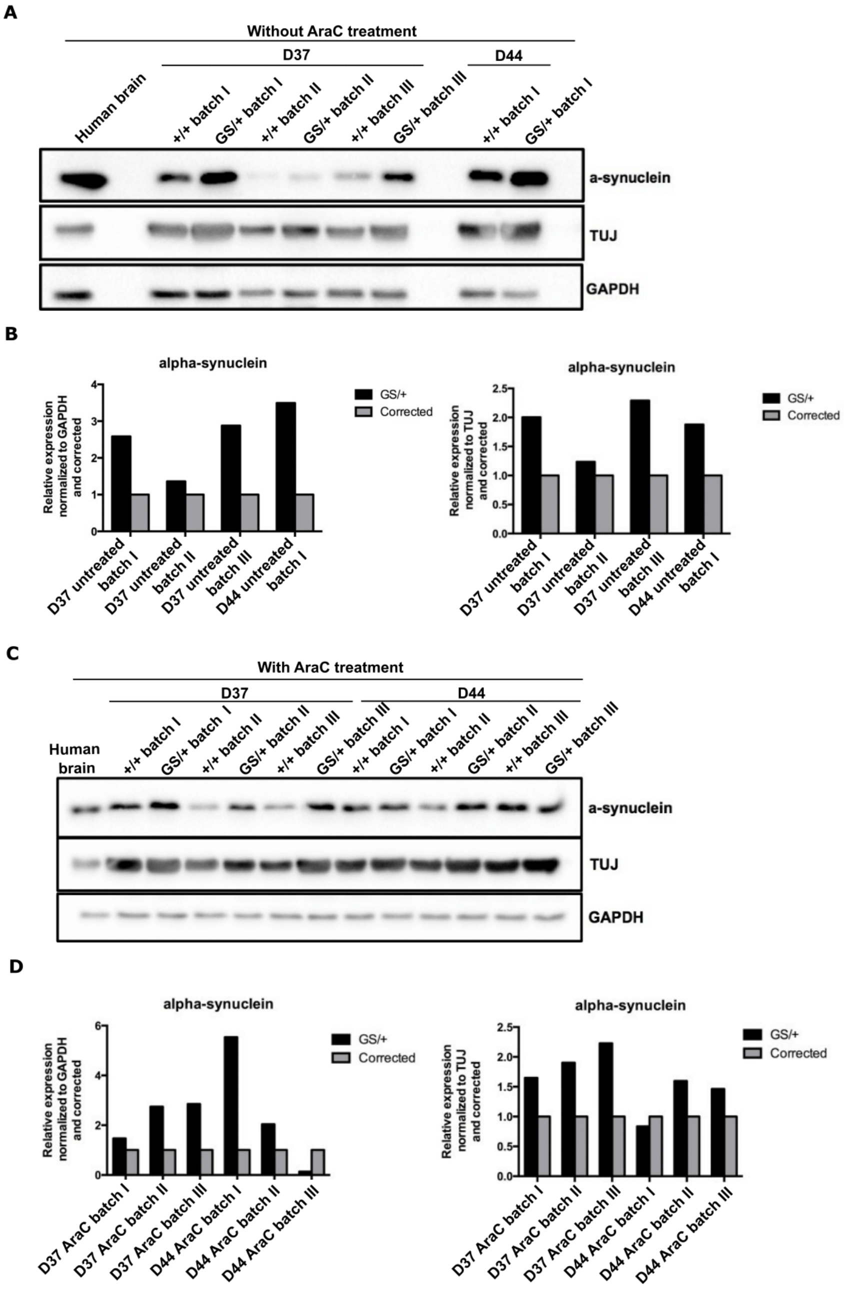


**Figure S2 (referring to Figure 1): LRRK2 G2019S mDA neurons contain higher levels of α-synuclein protein than isogenic control mDA neurons.** (**A**) D30 mDA neurons were cryopreserved, thawed and cultured for either 7 days (D37) or 14 days (D44) and lysed. α-synuclein, β-tubulin III (TUJ antibody) and GAPDH were detected by Western blotting. (**B**) Western blots were analyzed densiometrically and the α-synuclein level was expressed relative to GAPDH (left panel) or TUJ signal (right panel). (**C**) Experiment performed as in (A), but cultured neurons were treated with the cytostatic drug cytosine arabinoside (AraC) in order to prevent glial proliferation. (**D**) Analysis was performed as in (**B**).

### Figure S3

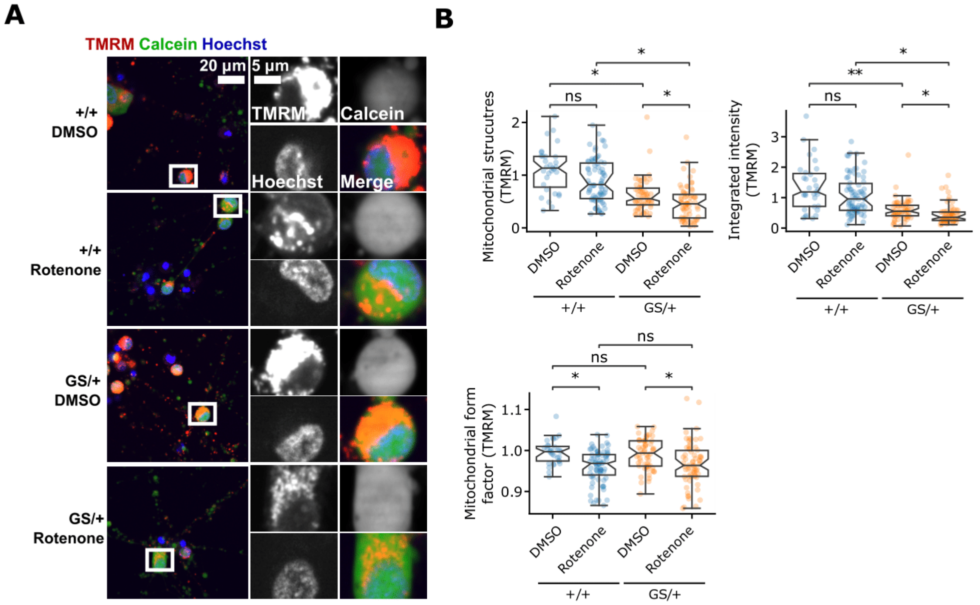


**Figure S3 (referring to Figure 1): LRRK2 G2019S mDA neurons are more sensitive to the mitochondrial stressor Rotenone.** (**A**) Representative images of cryopreserved D30 mDA neurons cultured for 7 days and treated with DMSO or Rotenone during the last 24 hours. Cells were stained with Hoechst, Calcein and TMRM and imaged. (**B**) Multiple mitochondrial features were quantified based on the TMRM stain including mitochondrial number, intensity, and shape.

### Figure S4

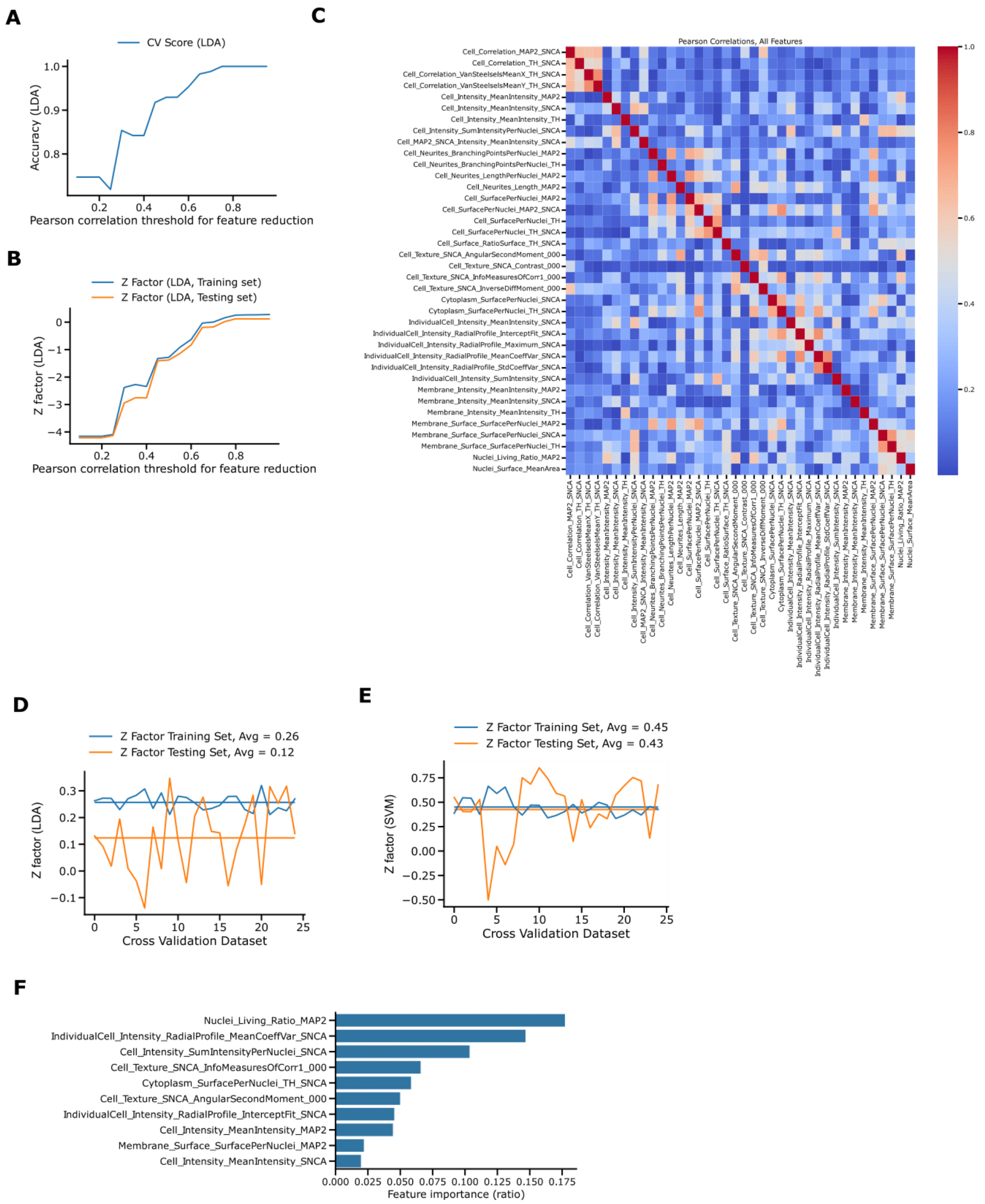


**Figure S4 (referring to Figure 3B): Feature selection, training and testing of Linear Discriminant Analysis (LDA) and Support Vector Machine (SVM) classifiers using Hoechst/TH/α-synuclein/MAP2 staining derived features originating from LRRK2 G2019S (GS/+) and isogenic control (+/+) neurons.** (**A**) LDA classification was used to select image features based on Hoechst/TH/α-synuclein/MAP2 staining that are not strongly correlated. LDA classifier accuracy is shown as a function of Pearson's correlation thresholds. (**B**) LDA training and testing set Z-factors between +/+ and GS/+ reference classes as a function of Pearson's correlation thresholds used to exclude correlated image features. (**C**) Pearson's correlation matrix of all selected image features for model training with threshold 0.80. (**D**) Performance of LDA classifier during 25 cycles of training (80% of data) and testing (20% of data) using 25 shuffled data sets. Z-factor values were calculated between reference classes +/+ and GS/+. (**E**) Same as in (**D**) but using SVM classification. (**D**) Ratios of feature contributions to Light Gradient Boosting Machine (LightGBM) classification of reference classes +/+ and GS/+.

### Figure S5

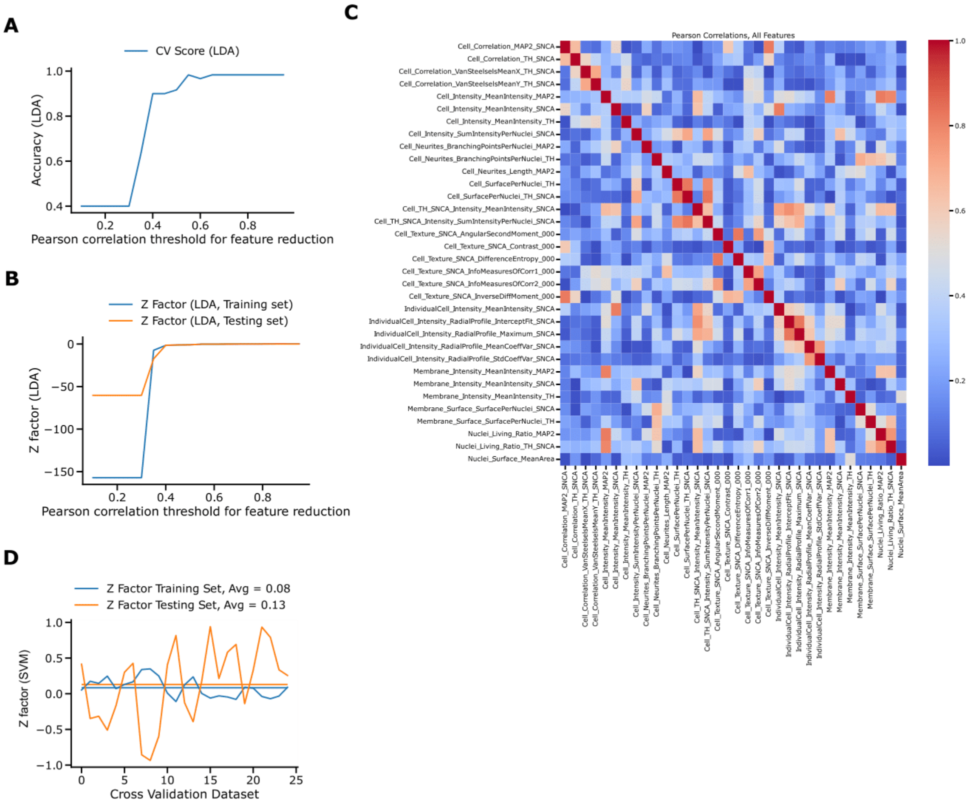


**Figure S5 (referring to Figure 3D):** Feature selection, training and testing of Support Vector Machine (SVM) classifier using Hoechst/TH/α-synuclein/MAP2 staining derived features originating from LRRK2 G2019S (GS/+) and isogenic control (+/+) neurons. (**A**) Linear Discriminant Analysis (LDA) classification was used to select image features based on Hoechst/TH/α-synuclein/MAP2 staining that are not strongly correlated. LDA classifier accuracy is shown as a function of Pearson's correlation thresholds. (**B**) LDA training and testing set Z-factors between +/+ and GS/+ reference classes as a function of Pearson's correlation thresholds used to exclude correlated image features. (**C**) Pearson's correlation matrix of all selected image features for model training with threshold 0.85. (D) Performance of SVM classifier during 25 cycles of training (80% of data) and testing (20% of data) using 25 shuffled data sets. Z-factor values were calculated between reference classes +/+ and GS/+.

### Figure S6

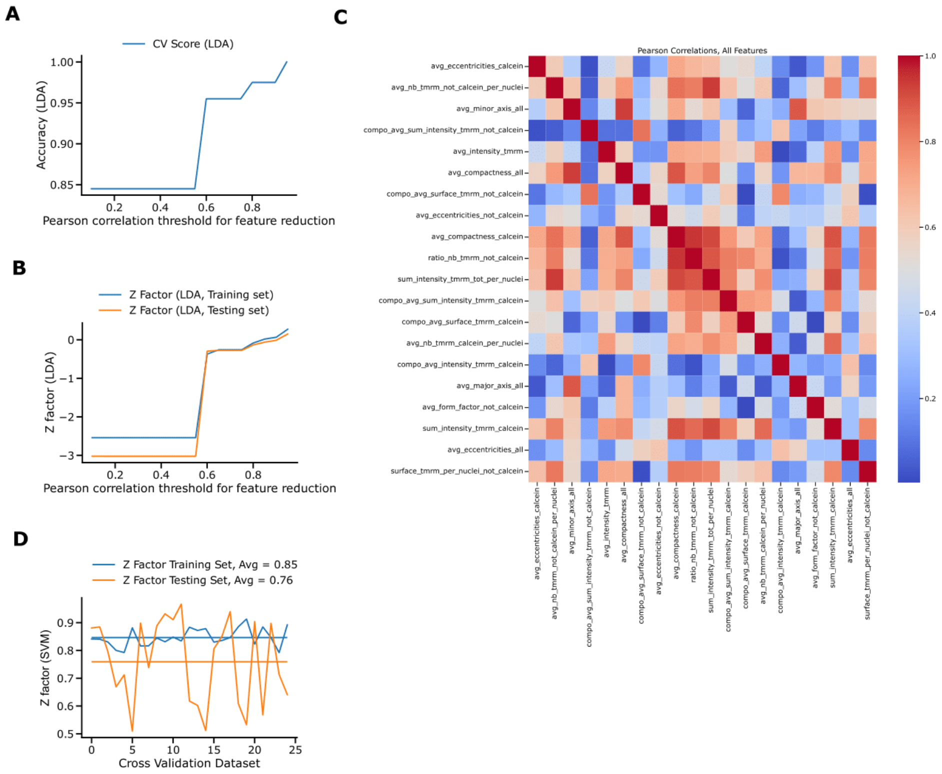


**Figure S6 (referring to Figure 3E): Feature selection, training and testing of Support Vector Machine (SVM) classifier using Hoechst/Calcein/TMRM staining derived features originating from LRRK2 G2019S (GS/+) and isogenic control (+/+) neurons.** (**A**) Linear Discriminant Analysis (LDA) classification was used to select image features based on Hoechst/Calcein/TMRM staining that are not strongly correlated. LDA classifier accuracy is shown as a function of Pearson's correlation thresholds. (**B**) LDA training and testing set Z-factors between +/+ and GS/+ reference classes as a function of Pearson's correlation thresholds used to exclude correlated image features. (**C**) Pearson's correlation matrix of all selected image features for model training with threshold 0.95. (**D**) Performance of SVM classifier during 25 cycles of training (80% of data) and testing (20% of data) using 25 shuffled data sets. Z-factor values were calculated between reference classes +/+ and GS/+.

### Figure S7

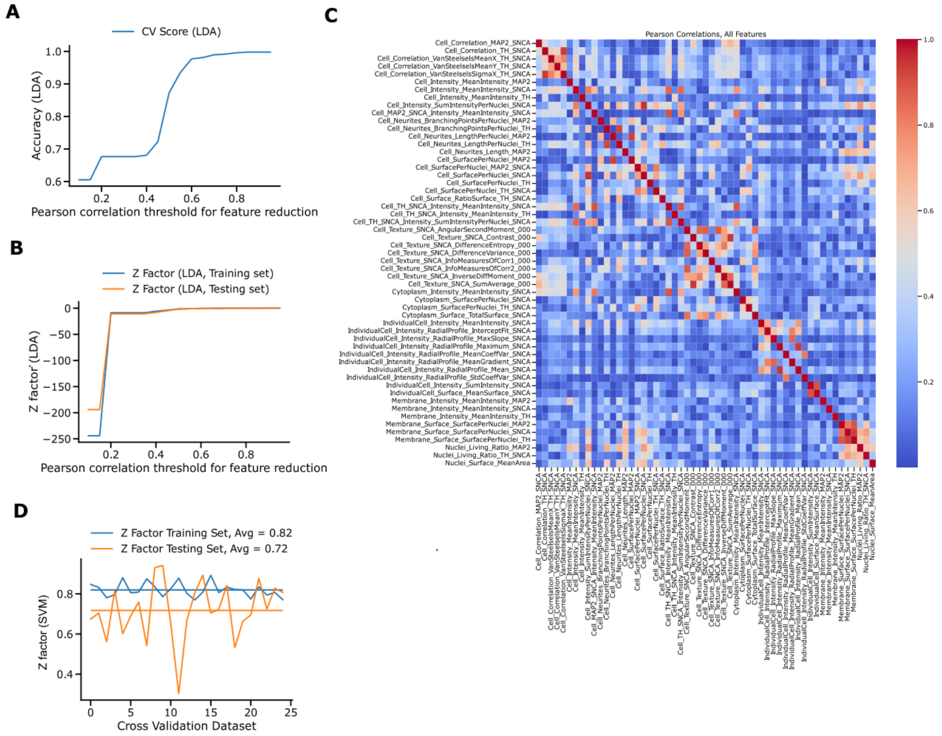


**Figure S7 (referring to Figure 4): Feature selection, training and testing of Support Vector Machine (SVM) classifier using Hoechst/TH/α-synuclein/MAP2 staining derived features originating from LRRK2 G2019S (GS/+) and isogenic control (+/+) neurons.** (**A**) Linear Discriminant Analysis (LDA) classification was used to select image features based on Hoechst/TH/α-synuclein/ MAP2 staining that are not strongly correlated. LDA classifier accuracy is shown as a function of Pearson's correlation thresholds. (**B**) LDA training and testing set Z-factors between +/+ and GS/+ reference classes as a function of Pearson's correlation thresholds used to exclude correlated image features. (**C**) Pearson's correlation matrix of all selected image features for model training with threshold 0.95. (**D**) Performance of SVM classifier during 25 cycles of training (80% of data) and testing (20% of data) using 25 shuffled data sets. Z-factor values were calculated between reference classes +/+ and GS/+.

### Figure S8

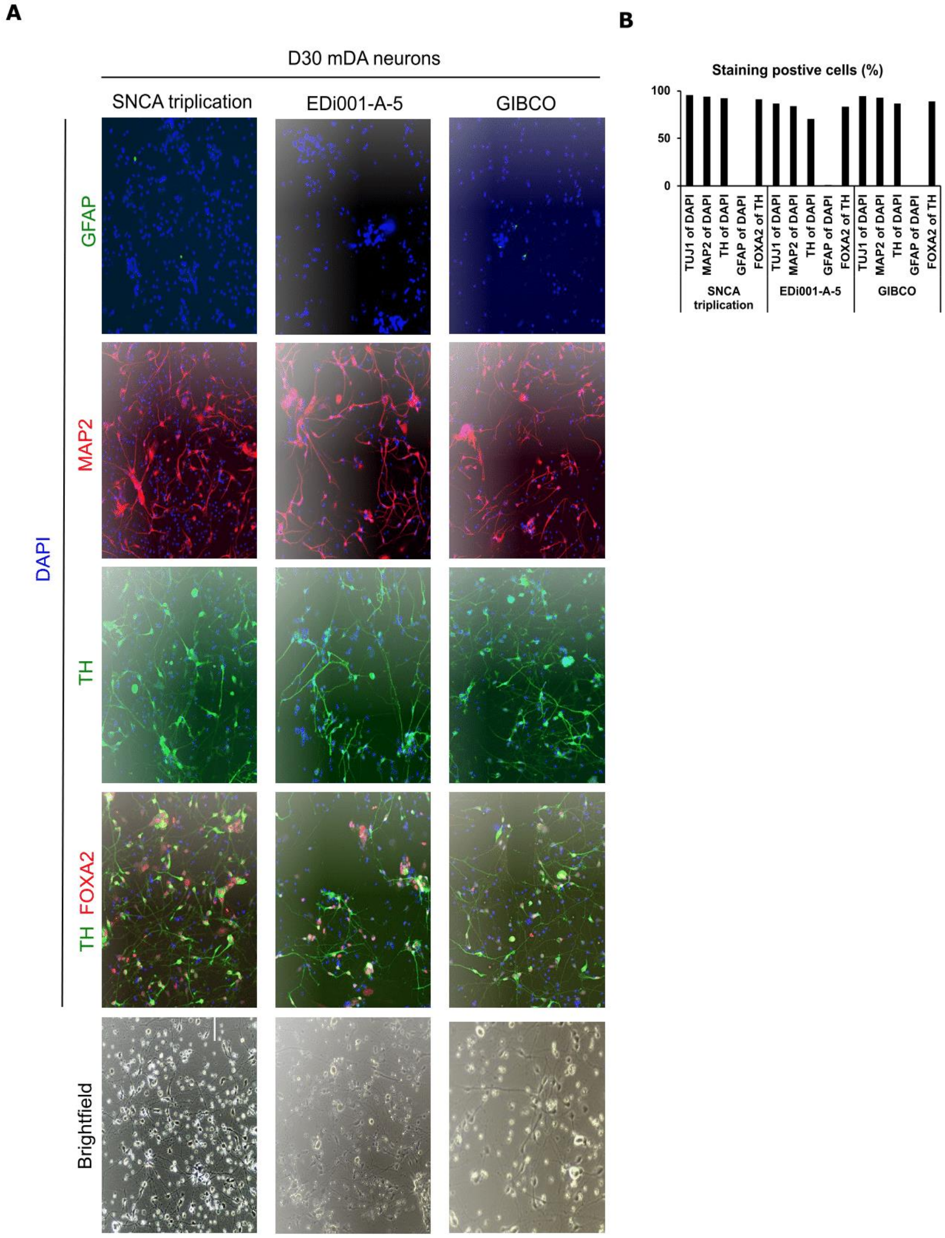


**Figure S8 (referring to Figure 5): Characterization of additional (non-LRRK2) human iPSC-derived mDA neurons.** (**A**) Batches of SNCA triplication, isogenic control (Edi001-A-5) and a genetically unrelated control (Gibco) iPSCs were differentiated into mDA neurons for 30 days and cryo-preserved at 2x106 cells per well. Thawed neurons were cultured for additional 7 days and stained for Tyrosine hydroxylase (TH), Forkhead box protein A2 (FOXA2), Microtubule-associated protein 2 (MAP2), and Glial Fibrillary Acidic Protein (GFAP). Representative images are shown. (**B**) Quantification of staining positive cells.

### Figure S9

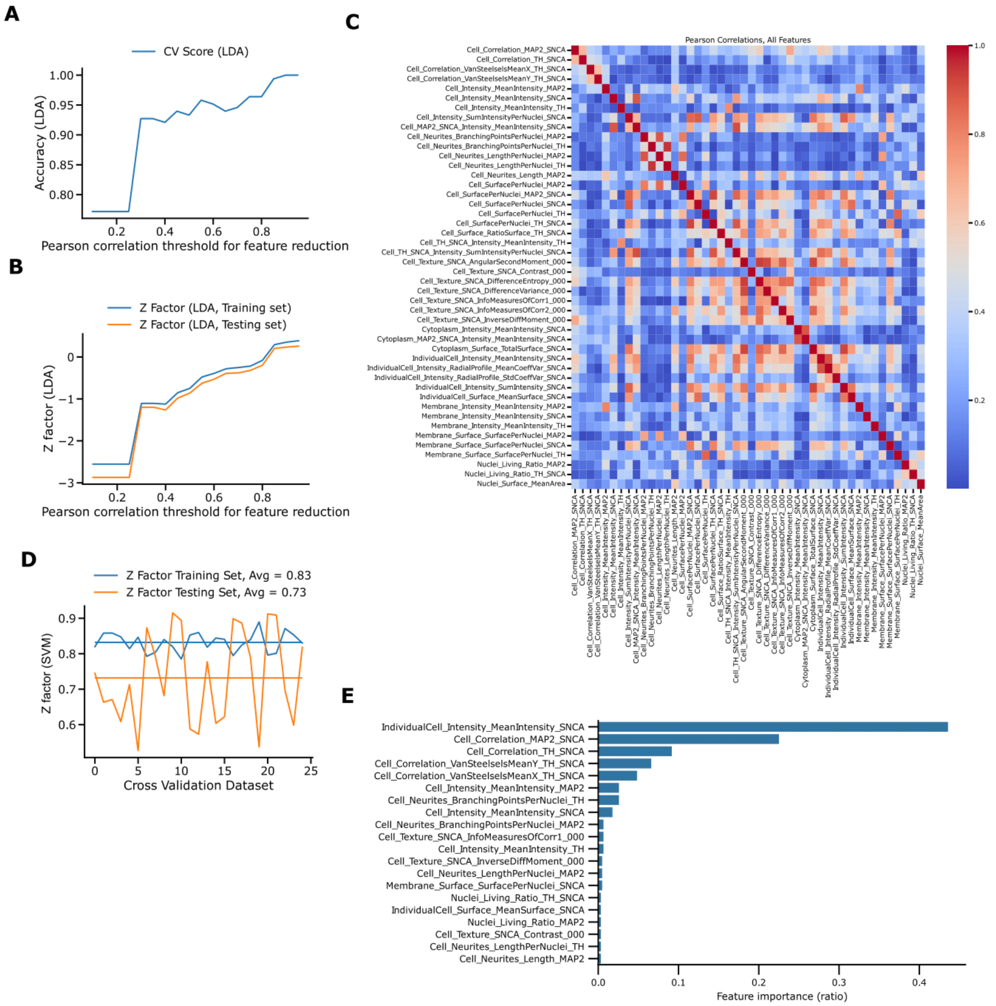


**Figure S9 (referring to Figure 5): Feature selection, training and testing of Support Vector Machine (SVM) classifier using Hoechst/TH/α-synuclein/MAP2 staining derived features originating from SNCA triplication and isogenic control neurons.** (**A**) Linear Discriminant Analysis (LDA) classification was used to select image features based on Hoechst/TH/α-synuclein/MAP2 staining that are not strongly correlated. LDA classifier accuracy is shown as a function of Pearson's correlation thresholds. (**B**) LDA training and testing set Z-factors between SNCA triplication and isogenic control reference classes as a function of Pearson's correlation thresholds used to exclude correlated image features. (**C**) Pearson's correlation matrix of all selected image features for model training with threshold 0.95. (**D**) Performance of SVM classifier during 25 cycles of training (80% of data) and testing (20% of data) using 25 shuffled data sets. Z-factor values were calculated between SNCA triplication and isogenic control reference classes. (**E**) Ratios of feature contributions to Light Gradient Boosting Machine (LightGBM) classification of reference classes SNCA triplication and isogenic control.

### Table S1

**Table S1: Description of extracted image features after Hoechst/TH/α-synuclein/MAP2 staining and fluorescence channel segmentation.**

| | Feature name | Feature description |
|---|---|---|
| 1 | Cell_Correlation_MAP2_SNCA | Pearson correlation between MAP2 and SNCA channel |
| 2 | Cell_Correlation_TH_SNCA | Pearson correlation between TH and SNCA channel |
| 3 | Cell_Correlation_VanSteelselsMeanX_TH_SNCA | Van Steelsel's cross correlation between TH and SNCA channel, shift on x-axis |
| 4 | Cell_Correlation_VanSteelselsMeanY_TH_SNCA | Van Steelsel's cross correlation between TH and SNCA channel, shift on y-axis |
| 5 | Cell_Correlation_VanSteelselsSigmaX_TH_SNCA | SD of Van Steelsel's cross correlation between TH and SNCA channel, shift on x-axis |
| 6 | Cell_Correlation_VanSteelselsSigmaY_TH_SNCA | SD of Van Steelsel's cross correlation between TH and SNCA channel, shift on y-axis |
| 7 | Cell_Intensity_MeanIntensity_MAP2 | Mean pixel intensity of MAP2 channel |
| 8 | Cell_Intensity_MeanIntensity_SNCA | Mean pixel intensity of SNCA channel |
| 9 | Cell_Intensity_MeanIntensity_TH | Mean pixel intensity of TH channel |
| 10 | Cell_Intensity_SumIntensityPerNuclei_SNCA | Integrated pixel intensity of SNCA channel normalized to number of nuclei |
| 11 | Cell_MAP2_SNCA_Intensity_MeanIntensity_SNCA | Mean pixel intensity of SNCA channel colocalized to MAP2 channel |
| 12 | Cell_Neurites_BranchingPointsPerNuclei_MAP2 | Dendritic branching points of MAP2 channel normalized to number of nuclei |
| 13 | Cell_Neurites_BranchingPointsPerNuclei_TH | Dendritic branching points of TH channel normalized to number of nuclei |
| 14 | Cell_Neurites_LengthPerNuclei_MAP2 | Dentritic network length of MAP2 channel normalized to number of nuclei |
| 15 | Cell_Neurites_LengthPerNuclei_TH | Dentritic network length of TH channel normalized to number of nuclei |
| 16 | Cell_Neurites_Length_MAP2 | Dentritic network length of MAP2 channel |
| 17 | Cell_Neurites_Length_TH | Dentritic network length of TH channel |
| 18 | Cell_SurfacePerNuclei_MAP2 | Surface pixels occupied by MAP2 channel, normalized to number of nuclei |
| 19 | Cell_SurfacePerNuclei_MAP2_SNCA | Surface pixels occupied by colocalized MAP2 and SNCA channel normalized to number of nuclei |
| 20 | Cell_SurfacePerNuclei_SNCA | Surface pixels occupied by SNCA channel, normalized to number of nuclei |
| 21 | Cell_SurfacePerNuclei_TH | Surface pixels occupied by TH channel, normalized to number of nuclei |
| 22 | Cell_SurfacePerNuclei_TH_SNCA | Surface pixels occupied by colocalized TH and SNCA channel normalized to number of nuclei |
| 23 | Cell_Surface_RatioSurface_TH_SNCA | Surface ratio occupied by colocalized TH and SNCA channel |
| 24 | Cell_Surface_TotalSurface_MAP2 | Surface pixels occupied by MAP2 channel |
| 25 | Cell_Surface_TotalSurface_SNCA | Surface pixels occupied by SNCA channel |
| 26 | Cell_Surface_TotalSurface_SNCA_MAP2 | Surface pixels occupied by colocalized SNCA and MAP2 channel |
| 27 | Cell_Surface_TotalSurface_TH | Surface pixels occupied by TH channel |
| 28 | Cell_Surface_TotalSurface_TH_SNCA | Surface pixels occupied by colocalized TH and SNCA channel |
| 29 | Cell_TH_SNCA_Intensity_MeanIntensity_SNCA | Mean pixel intensity of SNCA channel colocalized to TH channel |
| 30 | Cell_TH_SNCA_Intensity_MeanIntensity_TH | Mean pixel intensity of TH channel colocalized to SNCA channel |
| 31 | Cell_TH_SNCA_Intensity_SumIntensityPerNuclei_SNCA | Integrated pixel intensity of SNCA channel colocalized to TH channel normalized to number of nuclei |
| 32 | Cell_Texture_SNCA_AngularSecondMoment_000 | Haralick uniformity of distribution of gray levels at 0 degree shift |
| 33 | Cell_Texture_SNCA_AngularSecondMoment_045 | Haralick uniformity of distribution of gray levels at 45 degree shift |
| 34 | Cell_Texture_SNCA_AngularSecondMoment_090 | Haralick uniformity of distribution of gray levels at 90 degree shift |
| 35 | Cell_Texture_SNCA_AngularSecondMoment_135 | Haralick uniformity of distribution of gray levels at 135 degree shift |
| 36 | Cell_Texture_SNCA_Contrast_000 | Haralick contrast of gray levels at 0 degree shift |
| 37 | Cell_Texture_SNCA_Contrast_045 | Haralick contrast of gray levels at 45 degree shift |
| 38 | Cell_Texture_SNCA_Contrast_090 | Haralick contrast of gray levels at 90 degree shift |
| 39 | Cell_Texture_SNCA_Contrast_135 | Haralick contrast of gray levels at 135 degree shift |
| 40 | Cell_Texture_SNCA_Correlation_000 | Haralick correlation of gray levels at 0 degree shift |
| 41 | Cell_Texture_SNCA_Correlation_045 | Haralick correlation of gray levels at 45 degree shift |
| 42 | Cell_Texture_SNCA_Correlation_090 | Haralick correlation of gray levels at 90 degree shift |
| 43 | Cell_Texture_SNCA_Correlation_135 | Haralick correlation of gray levels at 135 degree shift |
| 44 | Cell_Texture_SNCA_DifferenceEntropy_000 | Haralick difference of randomness of gray levels at 0 degree shift |
| 45 | Cell_Texture_SNCA_DifferenceEntropy_045 | Haralick difference of randomness of gray levels at 45 degree shift |
| 46 | Cell_Texture_SNCA_DifferenceEntropy_090 | Haralick difference of randomness of gray levels at 90 degree shift |
| 47 | Cell_Texture_SNCA_DifferenceEntropy_135 | Haralick difference of randomness of gray levels at 135 degree shift |
| 48 | Cell_Texture_SNCA_DifferenceVariance_000 | Haralick difference of variance of gray levels at 0 degree shift |
| 49 | Cell_Texture_SNCA_DifferenceVariance_045 | Haralick difference of variance randomness of gray levels at 45 degree shift |
| 50 | Cell_Texture_SNCA_DifferenceVariance_090 | Haralick difference of variance randomness of gray levels at 90 degree shift |
| 51 | Cell_Texture_SNCA_DifferenceVariance_135 | Haralick difference of variance randomness of gray levels at 135 degree shift |
| 52 | Cell_Texture_SNCA_Entropy_000 | Haralick randomness of gray levels at 0 degree shift |
| 53 | Cell_Texture_SNCA_Entropy_045 | Haralick randomness of gray levels at 45 degree shift |
| 54 | Cell_Texture_SNCA_Entropy_090 | Haralick randomness of gray levels at 90 degree shift |
| 55 | Cell_Texture_SNCA_Entropy_135 | Haralick randomness of gray levels at 135 degree shift |
| 56 | Cell_Texture_SNCA_InfoMeasuresOfCorr1_000 | Haralick information measure of correlation 1 of gray levels at 0 degree shift |
| 57 | Cell_Texture_SNCA_InfoMeasuresOfCorr1_045 | Haralick information measure of correlation 1 of gray levels at 45 degree shift |
| 58 | Cell_Texture_SNCA_InfoMeasuresOfCorr1_090 | Haralick information measure of correlation 1 of gray levels at 90 degree shift |
| 59 | Cell_Texture_SNCA_InfoMeasuresOfCorr1_135 | Haralick information measure of correlation 1 of gray levels at 135 degree shift |
| 60 | Cell_Texture_SNCA_InfoMeasuresOfCorr2_000 | Haralick information measure of correlation 2 of gray levels at 0 degree shift |
| 61 | Cell_Texture_SNCA_InfoMeasuresOfCorr2_045 | Haralick information measure of correlation 2 of gray levels at 45 degree shift |
| 62 | Cell_Texture_SNCA_InfoMeasuresOfCorr2_090 | Haralick information measure of correlation 2 of gray levels at 90 degree shift |
| 63 | Cell_Texture_SNCA_InfoMeasuresOfCorr2_135 | Haralick information measure of correlation 2 of gray levels at 135 degree shift |
| 64 | Cell_Texture_SNCA_InverseDiffMoment_000 | Haralick homogeneity of gray levels at 0 degree shift |
| 65 | Cell_Texture_SNCA_InverseDiffMoment_045 | Haralick homogeneity of gray levels at 45 degree shift |
| 66 | Cell_Texture_SNCA_InverseDiffMoment_090 | Haralick homogeneity of gray levels at 90 degree shift |
| 67 | Cell_Texture_SNCA_InverseDiffMoment_135 | Haralick homogeneity of gray levels at 135 degree shift |
| 68 | Cell_Texture_SNCA_SumAverage_000 | Haralick sum of averages of gray levels at 0 degree shift |
| 69 | Cell_Texture_SNCA_SumAverage_045 | Haralick sum of averages of gray levels at 45 degree shift |
| 70 | Cell_Texture_SNCA_SumAverage_090 | Haralick sum of averages of gray levels at 90 degree shift |
| 71 | Cell_Texture_SNCA_SumAverage_135 | Haralick sum of averages of gray levels at 135 degree shift |
| 72 | Cell_Texture_SNCA_SumEntropy_000 | Haralick sum of gray level randomness at 0 degree shift |
| 73 | Cell_Texture_SNCA_SumEntropy_045 | Haralick sum of gray level randomness at 45 degree shift |
| 74 | Cell_Texture_SNCA_SumEntropy_090 | Haralick sum of gray level randomness at 90 degree shift |
| 75 | Cell_Texture_SNCA_SumEntropy_135 | Haralick sum of gray level randomness at 135 degree shift |
| 76 | Cell_Texture_SNCA_SumOfSquares_000 | Haralick sum of square gray level variance at 0 degree shift |
| 77 | Cell_Texture_SNCA_SumOfSquares_045 | Haralick sum of square gray level variance at 45 degree shift |
| 78 | Cell_Texture_SNCA_SumOfSquares_090 | Haralick sum of square gray level variance at 90 degree shift |
| 79 | Cell_Texture_SNCA_SumOfSquares_135 | Haralick sum of square gray level variance at 135 degree shift |
| 80 | Cell_Texture_SNCA_SumVariance_000 | Haralick sum of gray level variance at 0 degree shift |

**Table S1 (continued)**

| | | |
|---|---|---|
| 81 | Cell_Texture_SNCA_SumVariance_045 | Haralick sum of gray level variance at 45 degree shift |
| 82 | Cell_Texture_SNCA_SumVariance_090 | Haralick sum of gray level variance at 90 degree shift |
| 83 | Cell_Texture_SNCA_SumVariance_135 | Haralick sum of gray level variance at 135 degree shift |
| 84 | Cytoplasm_Intensity_MeanIntensity_SNCA | Mean pixel intensity of cytoplasmic SNCA channel |
| 85 | Cytoplasm_MAP2_SNCA_Intensity_MeanIntensity_SNC | Mean pixel intensity of cytoplasmic SNCA channel colocalized to MAP2 channel |
| 86 | Cytoplasm_SurfacePerNuclei_SNCA | Surface pixels occupied by cytoplasmic SNCA channel normalized to number of nuclei |
| 87 | Cytoplasm_SurfacePerNuclei_TH_SNCA | Surface pixels occupied by colocalized cytoplasmic TH and SNCA channel normalized to number of nuclei |
| 88 | Cytoplasm_Surface_TotalSurface_SNCA | Surface pixels occupied by cytoplasmic SNCA channel |
| 89 | IndividualCell_Intensity_MeanIntensity_SNCA | Mean pixel intensity of SNCA channel based on all individually segmented cells |
| 90 | IndividualCell_Intensity_RadialProfile_InterceptFit_SNCA | Fitted intercept of SNCA channel decay from center to edge |
| 91 | IndividualCell_Intensity_RadialProfile_MaxSlope_SNCA | Maximum steepness of SNCA channel decay from center to edge |
| 92 | IndividualCell_Intensity_RadialProfile_Maximum_SNCA | Maximum intensity of SNCA channel from center to edge |
| 93 | IndividualCell_Intensity_RadialProfile_MeanCoeffVar_SN | Mean SNCA channel dispersion from center to edge |
| 94 | IndividualCell_Intensity_RadialProfile_MeanGradient_SN | Mean shape of SNCA channel decay from center to edge |
| 95 | IndividualCell_Intensity_RadialProfile_Mean_SNCA | Mean intensity of SNCA channel from center to edge |
| 96 | IndividualCell_Intensity_RadialProfile_Median_SNCA | Median intensity of SNCA channel from center to edge |
| 97 | IndividualCell_Intensity_RadialProfile_Minimum_SNCA | Minimum intensity of SNCA channel from center to edge |
| 98 | IndividualCell_Intensity_RadialProfile_Q1_SNCA | First quartile intensity of SNCA channel from center to edge |
| 99 | IndividualCell_Intensity_RadialProfile_Q3_SNCA | Third quartile intensity of SNCA channel from center to edge |
| 100 | IndividualCell_Intensity_RadialProfile_SlopeFit_SNCA | Fitted slope of SNCA channel decay from center to edge |
| 101 | IndividualCell_Intensity_RadialProfile_StdCoeffVar_SNC | SD of SNCA channel dispersion from center to edge |
| 102 | IndividualCell_Intensity_RadialProfile_Std_SNCA | SD of SNCA channel intensity from center to edge |
| 103 | IndividualCell_Intensity_SumIntensity_SNCA | Integrated pixel intensity of SNCA channel based on all individually segmented cells |
| 104 | IndividualCell_Surface_MeanSurface_SNCA | Mean surface pixels occupied by SNCA channel based on all individually segmented cells |
| 105 | Membrane_Intensity_MeanIntensity_MAP2 | Mean pixel intensity of MAP2 channel on cellular edge |
| 106 | Membrane_Intensity_MeanIntensity_SNCA | Mean pixel intensity of TH channel on cellular edge |
| 107 | Membrane_Intensity_MeanIntensity_TH | Mean pixel intensity of MAP2 channel on cellular edge |
| 108 | Membrane_Surface_SurfacePerNuclei_MAP2 | Surface pixels on cellular edge occupied by MAP2 channel normalized to number of nuclei |
| 109 | Membrane_Surface_SurfacePerNuclei_SNCA | Surface pixels on cellular edge occupied by SNCA channel normalized to number of nuclei |
| 110 | Membrane_Surface_SurfacePerNuclei_TH | Surface pixels on cellular edge occupied by TH channel normalized to number of nuclei |
| 111 | Nuclei_Living_Ratio_MAP2 | Ratio of MAP2 channel positive nuclei |
| 112 | Nuclei_Living_Ratio_MAP2_SNCA | Ratio of MAP2 and SNCA channel positive nuclei |
| 113 | Nuclei_Living_Ratio_SNCA | Ratio of SNCA channel positive nuclei |
| 114 | Nuclei_Living_Ratio_TH | Ratio of TH channel positive nuclei |
| 115 | Nuclei_Living_Ratio_TH_SNCA | Ratio of TH and SNCA channel positive nuclei |
| 116 | Nuclei_Number_Big | Number of large nuclei |
| 117 | Nuclei_Number_Dead | Number of condensed/bright nuclei |
| 118 | Nuclei_Number_Living | Number of nuclei based on Hoechst channel |
| 119 | Nuclei_Number_MAP2 | Number of MAP2 channel positive nuclei |
| 120 | Nuclei_Number_MAP2_SNCA | Number of MAP2 and SNCA channel positive nuclei |
| 121 | Nuclei_Number_SNCA | Number of SNCA channel positive nuclei |
| 122 | Nuclei_Number_TH | Number of TH channel positive nuclei |
| 123 | Nuclei_Number_TH_SNCA | Number of TH and SNCA channel positive nuclei |
| 124 | Nuclei_Ratio_Dead | Ratio of condensed/bright nuclei |
| 125 | Nuclei_Ratio_Living | Ratio of nuclei not considered condensed/bright |
| 126 | Nuclei_Surface_MeanArea | Mean surface pixels of Hoechst channel |

#### Table S2

**Table S2: Description of extracted image features after Hoechst/Calcein/TMRM staining and fluorescence channel segmentation.**

| | Feature name | Feature description |
|---|---|---|
| 1 | avg_compactness_all | Average Compactness = area * 4 * pi / major axis : all the components |
| 2 | avg_compactness_calcein | Average Compactness = area * 4 * pi / major axis : only the components in the calcein mask |
| 3 | avg_compactness_not_calcein | Average Compactness = area * 4 * pi / major axis : only the components not in the calcein mask |
| 4 | avg_eccentricities_all | Average Eccentricity = sqrt(1 - (minor axis/major axis)^2) : all the components |
| 5 | avg_eccentricities_not_calcein | Average Eccentricity = sqrt(1 - (minor axis/major axis)^2) : only the components not in the calcein mask |
| 6 | avg_form_factor_all | Average Form Factor = 4*pi*area/perimeter : all the components |
| 7 | avg_form_factor_calcein | Average Form Factor = 4*pi*area/perimeter : only the components in the calcein mask |
| 8 | avg_form_factor_not_calcein | Average Form Factor = 4*pi*area/perimeter : only the components not in the calcein mask |
| 9 | avg_intensity_calcein | Average intensity of the calcein channel |
| 10 | avg_intensity_tmrm | Average intensity of the TMRM channel |
| 11 | avg_major_axis_all | Average major axis : all the components |
| 12 | avg_major_axis_calcein | Average major axis : only the components in the calcein mask |
| 13 | avg_major_axis_not_calcein | Average major axis : only the components not in the calcein mask |
| 14 | avg_minor_axis_all | Average minor axis : all the components |
| 15 | avg_minor_axis_calcein | Average minor axis : only the components in the calcein mask |
| 16 | avg_minor_axis_not_calcein | Average minor axis : only the components not in the calcein mask |
| 17 | avg_nb_tmrm_calcein_per_nuclei | Average number of components tmrm in calcein mask normalized per nulcei TMRM positive |
| 18 | avg_nb_tmrm_per_nuclei | Total number of components tmrm normalized per nulcei TMRM positive |
| 19 | avg_perimeters_all | Average perimeter of components : all the components |
| 20 | avg_perimeters_calcein | Average perimeter of components : only the components in the calcein mask |
| 21 | avg_perimeters_not_calcein | Average perimeter of components : only the components not in the calcein mask |
| 22 | compo_avg_intensity_tmrm_calcein | Average TMRM intensity per component : only the components in the calcein mask |
| 23 | compo_avg_intensity_tmrm_not_calcein | Average TMRM intensity per component : only the components not in the calcein mask |
| 24 | compo_avg_sum_intensity_tmrm_calcein | Average sum of TMRM intensity per component : only the components in the calcein mask |
| 25 | compo_avg_sum_intensity_tmrm_not_calcein | Average sum of TMRM intensity per component : only the components not in the calcein mask |
| 26 | compo_avg_surface_tmrm_calcein | Average surface of TMRM components : only the components in the calcein mask |
| 27 | compo_avg_surface_tmrm_not_calcein | Average surface of TMRM components : only the components not in the calcein mask |
| 28 | dead_nuclei | Number of dead cells |
| 29 | living_nuclei | Number of living cells |
| 30 | nuclei_tot | Total number of cells |
| 31 | ratio_dead_nuclei | Ratio of dead cells |
| 32 | ratio_living_nuclei | Ratio of living cells |
| 33 | ratio_nb_tmrm_calcein | Ratio of TMRM components in the calcein mask |
| 34 | ratio_nb_tmrm_not_calcein | Ratio of TMRM components in the not calcein mask |
| 35 | sum_intensity_calcein_per_nuclei | Sum of intensities of calcein channel normalized by living cells |
| 36 | sum_intensity_calcein_tot | Sum of intensities of calcein channel |
| 37 | sum_intensity_calcein_tot_per_nuclei | Sum of intensities of calcein channel normalized by living cells |
| 38 | sum_intensity_tmrm_calcein | Sum of intensities of TMRM channel in the calcein mask |
| 39 | sum_intensity_tmrm_calcein_per_nuclei | Sum of intensities of TMRM channel in the calcein mask normalized by calcein cell positive cell number |
| 40 | sum_intensity_tmrm_not_calcein | Sum of intensities of TMRM channel not in the calcein mask |
| 41 | sum_intensity_tmrm_not_calcein_per_nuclei | Sum of intensities of TMRM channel not in the calcein mask normalized by calcein cell positive cell number |
| 42 | sum_intensity_tmrm_tot | Sum of intensities of TMRM channel |
| 43 | sum_intensity_tmrm_tot_per_nuclei | Sum of intensities of TMRM channel normalized by living cell number |
| 44 | surface_calcein | Surface of calcein channel above the threshold |
| 45 | surface_calcein_per_nuclei_calcein | Surface of calcein channel above the threshold normalized by calcein positive cell number |
| 46 | surface_tmrm_calcein | Total surface of TMRM in calcein mask. |
| 47 | surface_tmrm_per_nuclei_not_calcein | Surface of TMRM not in the calcein mask normalized by negative calcein cell number |
| 48 | surface_tmrm_per_nuclie_calcein | Surface of TMRM in the calcein mask normalized by positive calcein cell number |
| 49 | total_nb_tmrm | Total number of components TMRM |
| 50 | total_nb_tmrm_calcein | Total number of components TMRM in calcein mask |
| 51 | avg_eccentricities_calcein | Average Eccentricity = sqrt(1 - (minor axis/major axis)^2) : only the components in the calcein mask |
| 52 | avg_nb_tmrm_not_calcein_per_nuclei | Number of TMRM components not in calcein mask normalized by negative calcein cell number |
| 53 | surface_tmrm | Total surface of TMRM channel above the threshold |
| 54 | total_nb_tmrm_not_calcein | Total number of TMRM components not in the calcein mask |

### Table S3

**Table S3: Used cell lines, antibodies, primers, and key reagents.**

| **Cell lines** | | | | | |
|---|---|---|---|---|---|
| **Genotype** | **hPSCreg name** | **Donor source** | **Provider** | **Reprogramming method** | **Ref** |
| LRRK2 G2019S | STBCi004-B (GS/+) | Female, dermal fibroblasts | Distributor: EBiSC; Generator: StemBANCC | Non-integrating Sendai virus | (Morrison et al., 2015) |
| Corrected LRRK2 G2019S mutation in STBCi004-B | STBCi004-B-1 (+/+) | Female, dermal fibroblasts | Distributor: EBiSC; Generator: StemBANCC | Non-integrating Sendai virus | (Morrison et al., 2015) |
| SNCA triplication | EDi001-A (AST23) | Female, dermal fibroblasts | Distributor: EBiSC; Generator: University of Edinburgh | Integrating Retro virus | (Devine et al., 2011; Gwinn et al., 2011) |
| Corrected SNCA triplication in EDi001-A | EDi001-A-5 (AST23-2KO-8B) | Female, dermal fibroblasts | Distributor: EBiSC; Generator: University of Edinburgh | Integrating Retro virus | (Devine et al., 2011; Gwinn et al., 2011) |
| No known mutations | TMOi001-A (Gibco A18944) | Female, CD34+ cord blood | Distributor: EBiSC; Generator: ThermoFisher | Non-integrating Epstein-Barr virus | (Burridge et al., 2011) |
| **Antibodies** | | | | | |
| **Antibody** | **Assay** | **Dilution** | **Distributor** | | |
| TH | ICC (quality control) | 1/1500, 5% BSA + PBS | Millipore, #AB152 | | |
| FOXA2 | ICC (quality control) | 1/200 5% BSA + PBS | Biotechne, #AF2400 | | |
| MAP2 | ICC (quality control) | 1/1000, 5% BSA + PBS | Sigma, #M4403 | | |
| GFAP | ICC (quality control) | 1/1500, 5% BSA + PBS | Merck, #AB5804 | | |
| α-synuclein | WB | 1/1000 in Invitrogen™ iBind™ Flex Solution Kit | Novus, #NBP1-05194 | | |
| TUBB3/TUJ | WB | 1/1000, 5% milk + TBS-T, 2h, RT | Cell Signaling, #5568 | | |
| LRRK2 | WB | 1/500, 5% milk + TBS-T, 2h, RT | NeuroMab, #N241A/34 | | |
| Actin | WB | 1/1000, 5% milk + TBS-T, 2h, RT | Cell Signaling, #58169 | | |
| GAPDH | WB | 1/1000 in Invitrogen™ iBind™ Flex Solution Kit | Cell Signaling, #2118 | | |
| Anti-Rabbit IgG HRP-linked | WB | 1/1000, 5% milk + TBS-T, 1h, RT | Cell Signaling, #7074S | | |
| α-synuclein rabbit | ICC | 1/500, 5% FBS + 0.1% Triton X-100 + PBS, overnight, 4°C | Abcam, #138501 | | |
| TH | ICC | 1/1000, 5% FBS + 0.1% Triton X-100 + PBS, overnight, 4°C | Merck, #T2928 | | |
| MAP2 | ICC | 1/5000, 5% FBS + 0.1% Triton X-100 + PBS, overnight, 4°C | Novus, #NB300-213 | | |
| pS129 α-synuclein | ICC | 1/500, 5% FBS + 0.1% Triton X-100 + PBS, overnight, 4°C | Cell Signaling, #23706S | | |
| α-synuclein mouse | ICC | 1/500, 5% FBS + 0.1% Triton X-100 + PBS, overnight, 4°C | BD Biosciences, #610787 | | |
| Alexa Fluo 488, Anti-Mouse | ICC | 1/1000, 5% FBS + 0.1% Triton X-100 + PBS, 1h, RT | ThermoFisher, #A11001 | | |
| Alexa Fluor 647 Anti-Chicken | ICC | 1/250, 5% FBS + 0.1% Triton X-100 + PBS, 1h, RT | Jackson Immuno Research, #703-605-155 | | |

**Table S3 (continued)**

| | | | | | |
|---|---|---|---|---|---|
| Alexa Fluo 555, Anti-Rabbit | ICC | 1/1000, 5% FBS + 0.1% Triton X-100 + PBS, 1h, RT | ThermoFisher, #A21429 | | |
| TMRM | ICC | 25nM, Differentiation medium, 30 min, 37°C | ThermoFisher, #T668 | | |
| Calcein | ICC | 1.25µM, Differentiation medium, 30 min, 37°C | ThermoFisher, #C3100MP | | |
| Hoechst 33342 | ICC | 1/2000, 30 min, 37°C for live staining or 1/3000 1h, RT for ICC | Sigma, # 14533 | | |
| **Primers** | | | | | |
| **LRRK2 Primer** | **Melting temp (˚C)** | **Annealing temp (˚C)** | **GC content (%)** | **Sequence** | **Product size (bp)** |
| FWD | 58.98 | 55.98 | 50 | GCTTGTTGTTGGAC AGCTGA | Genomic: 1262 |
| REV | 58.97 | 55.97 | 50 | GCTTGTTGTTGGAC AGCTGA | mRNA: 224 |
| **Media** | | | | | |
| **Reagents** | **Stock** | **Dilution** | **Distributor** | | |
| Basal medium | | | | | |
| Neurobasal medium | | 1 | Gibco, #21103-049 | | |
| GlutaMAX | 100x | 1:100 | Gibco, #25030-081 | | |
| Pen/Strep | 100x | 1:100 | Gibco, #15070-063 | | |
| B27 | 50x | 1:50 | Gibco, #12587-010 | | |
| Differentiation medium | | | | | |
| Basal medium | | 1 | | | |
| BDNF | 10 µg/mL | 1:500 | Cell Guidance Sys., #GFH1-10 | | |
| GDNF | 10 µg/mL | 1:500 | Cell Guidance Sys., #GFH2-10 | | |
| LAAP | 221 mM | 1:1000 | Sigma, #A8960-5G | | |
| DAPT | 10 mM | 1:1000 | Axon Medchem, #1484 | | |
| TGF-β3 | 1 µg/mL | 1:1000 | Peprotech, #100-36E | | |
| dbcAMP | 100 mM | 1:200 | Enzo, #BML-CN125-0100 | | |

#### Table S4

**Table S4: Machine learning (ML) summary table as per data, optimization, model and evaluation (DOME) recommendations (Walsh et al., 2021).**

| | | |
|---|---|---|
| **Data** | Provenance | Source of all used data where the experiments and image analysis described in this paper. **Tab. 2** contains information on all generated datasets. |
| | Data splits | Available data was split into 80% training data and 20% testing data. No separate validation set was used due to the use of k-fold cross validation. |
| | Redundancy between data splits | Train and test sets were generated using scikitlearn's train_test_split function with enabled stratification and random seeding to ensure that relative class frequencies were approximately preserved in each train and test set. |
| | Availability of data | All data is available in the **Supporting Material**. Data splits can be reproduced using the provided Jupyter Notebooks (link to GitHub in online version of paper). |
| **Optimization** | Algorithm | Established supervised binary classification algorithms were used. LDA was used for feature reduction and initial classification. Non-linear SVM proved to be more accurate and was subsequently used for classification tasks followed by leave-one-out (LOO) analysis to determine feature contributions. Tree-based LightGBM was used to confirm feature contributions determined by LOO. |
| | Meta-prediction | LDA was used during the feature selection process. See also "Features" section. |
| | Data encoding | Outliers were removed by applying a 3xSD window around each feature's median. Data was transformed on the same scale using the following formula: $(x_{Feature\_N} – median_{Feature\_N})/SD$. |
| | Parameters | SVM parameters were systematically identified using scikitlearn's GridSearchCV function. The parameters of all other models were determined empirically.<br>LDA: solver='eigen', n_components=1, shrinkage='auto'<br>SVM: probability=True, kernel='rbf', C=0.1-0.5 (depends on dataset, see provided Jupyter notebooks), gamma='scale'<br>LightGBM: boosting_type='goss', n_estimators=10000, class_weight = 'balanced' |
| | Features | **Tab. 2** contains information on all generated datasets including the total number of features. For feature reduction, Pearson correlation was used to exclude strongly correlated image features. For each dataset, LDA was used to determine a Pearson correlation cut-off value which maximized the accuracy of the classification and the Z factor between the two reference classes. The selected number and type of features are detailed per data set in the provided Jupyter notebooks. |
| | Fitting | Since the number of total features was high (>50) we used several approaches to prevent overfitting. GridSearchCV was used to optimize the SVM regularization parameter C. For LDA shrinkage was set to 'auto'. Additionally, we used k-fold cross validation to use a maximum of data for training and to prevent data leaking into the test set. |
| | Regularization | GridSearchCV was used to optimize the SVM regularization parameter C. The LDA parameter shrinkage was set to 'auto'. |
| | Availability of configuration | All configurations of all models for all data sets are reported in the provided Jupyter Notebooks (link to GitHub in online version of paper). |
| **Model** | Interpretability | Due to its non-linear nature the used SVM algorithm is not fully interpretable, but we performed LOO analysis to extract the approximate contribution of each used feature to the prediction. |
| | Output | SVM provides the classification probability to belong to one of the two used classes. |
| | Execution time | The entire workflow completes in ca. 80 seconds on a standard desktop computer. |
| | Availability of software | Jupyter Notebooks for each used data set are provided on GitHub. All dependencies are Python-based and freely available. |
| **Evaluation** | Evaluation method | 25-fold cross-validation was used to validate the SVM and LDA models for all datasets. |
| | Performance measures | F1 score, recall and accuracy are reported. |
| | Comparison | We did not benchmark our models with previously existing data, since appropriate mDA neuron data is not available. |
| | Confidence | F1 score, recall and accuracy are reported together with confidence intervals. |
| | Availability of evaluation | Model evaluation can be reproduced using the Jupyter Notebooks for each used data set provided on GitHub. |

#### Supplemental experimental procedures

##### Western blotting

Samples were extracted from two differentiation batches of GS/+ and +/+ mDA neurons. Cell lysis was performed in RIPA buffer. Human cortex lysate served as control. Samples were loaded in NuPAGE LDS Sample Buffer (4x) (ThermoFisher, #NP0007) and ran on NUPAGE Novex 3-8% Tris-acetate gels (ThermoFisher, #EA03752BOX) immersed in NUPAGE Tris-acetate SDS running buffer (1x) (ThermoFisher, #LA0041) for 1 hour at 150V. Gels were transferred to PVDF membranes (Biorad, #1620177) in transfer buffer (25mM Tris, 192mM Glycine, 10% Methanol (pre-cooled)) overnight at 60V and room temperature. Membrane blocking was performed in 5% skim milk powder (ThermoFisher, #LP0031B) in TBS-T for 1 hour at room temperature. Used primary and secondary antibodies are summarized in **Table S3**. The membrane was developed using Super Signal® West Femto Maximum Sensitivity Substrate (ThermoFisher, #34095).

##### RT-qPCR

Samples were extracted from two differentiation batches of GS/+ and +/+ mDA neurons. Sample extraction was performed using a Maxwell 16 total RNA purification kit and the Maxwell RSC instrument. RNA concentration was determined using a NanoDrop ™. Quality control was performed using an Agilent 2100 Bioanalyzer. All samples had a RNA integrity number (RIN) value of 10. cDNA synthesis was performed using the qScript cDNA synthesis kit (Quantabio, #95047) on 500 ng RNA. The cDNA was diluted 1:5 in $ddH_20$ and GoTaq SYBR green 2x Super Mix (Promega) was used on a ViiA 7 Real-Time PCR System (ThermoFisher) using the primers indicated in **Table S3**.
